# CDK1 and MAPK/Erk prime maternal proteins for Calcineurin-dependent dephosphorylation during the oocyte-to-embryo transition in *Drosophila*

**DOI:** 10.64898/2026.11.20.768146

**Authors:** Jonathon M. Thomalla, William J. Comstock, Steve Dorus, Mariana F. Wolfner

## Abstract

The transition from oocyte to embryo relies on the utilization of maternal macromolecules that accumulate during oogenesis. Change in phosphorylation status of maternal proteins is thought to drive developmental events during egg activation (EA) given that transcription does not occur during this time. To determine how mature oocytes acquire phosphorylation signatures on maternal proteins that are compatible with phospho-changes at EA, we analyzed phosphorylation dynamics during oocyte maturation (OM). We found widespread acquisition of phosphorylation at CDK1-MAPK sites during OM, and these sites were largely dephosphorylated during EA. We determined that the calcium-stimulated phosphatase Calcineurin is required for dephosphorylation of CDK1-MAPK substrates at EA; genetic perturbation of CDK1 or Calcineurin altered phosphorylation patterns at EA and induced precocious embryonic translation in oocytes. Ex vivo developmental assays revealed crosstalk between MAPK and CDK1 during OM: while depletion of either MAPK or CDK1 alone mildly impacts gross oocyte morphogenesis during OM, reducing the activity of both together dramatically reduced OM efficiency. Further, in MAPK-depleted conditions, CDK1 activity levels drastically increase in a temporally-sensitive manner, suggesting compensation for MAPK by CDK1 to ensure faithful establishment of maternal phospho-protein load. Finally, we show that the receipt of sperm during fertilization has no impact on phosphorylation events at EA. Our analyses connect the earliest events in embryonic initiation to the factors that keep oocytes arrested, providing novel insights as to the dynamics and mechanisms of maternal phospho-protein regulation during the transition from oogenesis to embryogenesis.

## Introduction

The developing egg cell, or oocyte, undergoes the terminal stages of its development without detectable transcription^[1, 2]^. Post-transcriptional regulation thus dictates the transition from oogenesis to embryogenesis. Egg activation (EA), or the oocyte-to-embryo transition, terminates oogenesis and initiates embryogenesis^[3]^. Prior studies have implicated protein phosphorylation changes as key regulators of EA^[4–18]^. However, it is unknown how a mature oocyte establishes a phosphorylation profile that is compatible with the initiation of embryo development or how phospho-changes elicit, maintain, and release developmental arrest.

EA is a broad transition affecting global cellular processes including meiotic resumption and completion, translational reprogramming and stored mRNA handling, and disposal of cellular factors essential for oocyte development and maintenance but detrimental to embryo development ^[3, 19]^. Successful EA is critical for animal development, and many of its features display deep conservation, as well as evolutionary flexibility^[9, 11, 20]^. Studies in model organisms, including the fruit fly *Drosophila melanogaster*, have identified EA genes with human orthologs, including regulatory genes, highlighting the utility of such organisms for the study of EA*^[4, 5]^*.

During *Drosophila* oogenesis, the oocyte grows in a modular assembly line of sequentially developing follicles, called egg chambers^[21, 22]^. Egg chambers are morphologically divided into discrete Stages denoted 1-14^[23]^, and transcription within oocytes is absent across most of these stages ^[24]^. At Stage 12, during the last 4-5 hours of oogenesis, the oocyte is released from Prophase I arrest and begins the process of Oocyte Maturation (OM)^[25]^. OM entails the release from Prophase I arrest through the establishment of Metaphase I arrest^[26, 27]^, and includes germinal vesicle (GV) breakdown and morphogenetic events like dorsal appendage construction. The fully mature Stage 14 oocyte remains arrested in Metaphase I until it leaves the ovary and encounters mechanical stress, which initiates EA by opening mechanically-gated calcium channels and inducing a cytosolic calcium rise shortly before the egg is fertilized in the female reproductive tract^[28, 29]^.

In *Drosophila*, the rise in calcium initiated at EA triggers phosphorylation changes that depend on the calcium-activated phosphatase Calcineurin. Calcineurin is essential for EA: prior analyses determined that eggs lacking Calcineurin do not begin embryogenesis and exhibit aberrant phosphorylation profiles^[5, 12]^. Meiosis and translation regulators are among Calcineurin’s downstream targets^[5]^. However, the enzymes that prepare a mature oocyte for these critical phosphorylation changes at EA are unknown. The cyclin dependent kinase CDK1 (cdc2) regulates metaphase arrest^[30]^, and its function depends partially on its interactions with different Cyclins^[31]^. CDK1 targets such as Greatwall and Endos contribute to release from metaphase arrest by activating PP2A^[18]^, which in turn activates the Anaphase Promoting Complex (APC/C) by dephosphorylating its cofactor Cdc20/Fzy^[32, 33]^. In other species, CDK1 regulates translation by phosphorylating regulators like 4EBP-1, and CDK1’s activity can be influenced by availability of Cyclin B^[34, 35]^. In Drosophila, CDK1 is thought to contribute to translational arrest by phosphorylating and inactivating the Pan Gu Kinase (PNG) subunit Gnu^[16, 36]^.

The gene products that regulate both OM and EA (and early embryo development) are spatiotemporally linked, warranting analysis of both transitions together rather than separately ^[4–6, 37–39]^. This study integrates quantitative phosphoproteomics and genetics to define the post-translational modifications and regulators that are relevant for both OM and EA in *Drosophila,* and determines that fertilization is not required for phospho-signaling induced by EA. During OM, the maternal proteome is subjected to phosphorylation by CDK1 and the mitogen activated protein kinase (MAPK) Erk, whose activity is partially counteracted by Calcineurin at EA. Genetic perturbation of CDK1 and Calcineurin revealed that these enzymes antagonize PNG-dependent translation initiation; reduction or overabundance of these factors, respectively, mistimes the oocyte-to-embryo transition. Together, this study identifies novel candidate effectors of OM and EA and defines synergistic requirements between key regulatory kinases and phosphatases for successful development.

## Methods

### Sample preparation for phospho-tandem mass spectrometry (MS/MS)

Stage 12 and Stage 14 oocytes were dissected from 3-7 day-old females reared at 29°C and fed on dry yeast. Ovaries were dissected in a hypertonic buffer lacking calcium (55mM NaAcetate, 40mM KAcetate, 1.2mM MgCl_2_, 110mM sucrose, 100mM HEPES) to prevent premature activation. Oocytes were freed from ovarioles by vigorous pipetting and samples were collected in <30-minute intervals before being transferred to Eppendorf tubes. Residual media was removed and samples were flash frozen in liquid nitrogen prior to storage at −80°C. Embryos and eggs were collected in <30-minute intervals on grape juice plates supplemented with yeast paste. Eggs/embryos were washed with egg wash buffer (0.4% NaCl, 0.03% Triton), transferred to Eppendorf tubes, and residual media was removed before flash freezing.

To generate protein lysates, samples were ground with pestles at 4°C in a phosphatase inhibitor buffer (62.5mM HEPES, 2% SDS, 1mM TPCK, 1mM TLCK, 1.5mM Benzamidine HCl, 2ug/mL aprotinin, 0l5ug/mL leupeptin, 0.7ug/mL pepstatin, 1mM sodium orthovanadate, 10mM beta-glycerophosphaste, 20mM sodium fluoride) and lysates were combined to yield approximately 500 oocytes per replicate. Lysates were centrifuged at 20,000g for 30 minutes at 4°C and supernatants were collected and flash frozen. For quality control, protein lysates were monitored for phospho-stability by western blotting for Gnu, which exists in a mostly phosphorylated form in the mature oocyte but is predominantly dephosphorylated in the activated egg (Fig. S1E). Gnu ran at a higher molecular weight in mature oocytes than in activated eggs, indicating maintenance of phospho-modifications in our samples.

### Protein digestion and tandem mass tag (TMT) labeling

Protein concentration for each sample was determined by running on 10% precast Bis-Tris SDS Page (Invitrogen) along with serial amounts of E. coli lysates serving as a standard curve. A total of 120 µg protein of each sample for 16 samples (10 included in this study, 6 analyzed in a separate study) was used for further processing in the following buffer: 50mM TEAB pH 8.5, 6M Urea, 2M Thiourea, 2% SDS.

Samples were reduced with 10 mM tris(2-carboxyethyl)phosphine for 1 hour at 34 °C, alkylated with 27 mM iodoacetamide for 1 hour in the dark and then quenched with a final concentration of 21 mM Dithiothreitol. Each sample was digested separately using the S-Trap Mini Spin column (Protifi, Huntington NY). After quenching, 12% phosphoric acid was added to a final concentration of 1.2%. Followed by 1:7 dilution (v/v) with 90% methanol, 0.1M TEAB pH 8.5. The samples were then placed into spin columns and centrifuged 4000g for 30 seconds, then washed three times with 400 µl 90% methanol, 0.1 M TEAB pH 8.5.

Digestion was performed with 125µl trypsin at 50 ng/µl (1:20 w/w) in 50 mM TEAB pH 8.5 added to the top of spin columns. Spin columns were incubated for 16 hours at 37 °C. Following incubation, the digested peptides were eluted off the S-trap column sequentially with 80µl each of 50 mM TEAB pH 8.5 followed by 0.2% formic acid, 50% acetonitrile, and 0.2% formic acid and dried. All dried samples were resuspended with 100 µl Optima water and re-dried to ensure the complete removal of formic acid. Immediately before labeling, each sample was reconstituted in 100 µl 0.1M TEAB pH 8.5. The TMT 16-plex labels (0.5 mg dried powder) were reconstituted with 50 µL of anhydrous ACN prior to labeling and added with 1: 2 ratio to each digested sample and incubated for 1 hour at room temperature. The labeled peptides were cleaned up by solid phase extraction on MCX Cartridges (Waters, Milford, MA). Incorporation of the TMT label was verified using Orbitrap Fusion (Thermo-Fisher Scientific, San Jose, CA). The eluted peptides were divided into four aliquots and evaporated. One aliquot containing 300 µg labeled peptides was used for the first dimensional LC fractionation via a high pH reverse phase chromatography, and three additional aliquots each containing 800 µg labeled peptides were used for enrichment of phosphopeptides as described below.

### High pH reverse phase (hpRP) fractionation

The hpRP chromatography was carried out using a Dionex UltiMate 3000 HPLC system (Thermo Scientific, Sunnyvale, CA). The TMT 16-plex tagged tryptic peptides were reconstituted in buffer A (20 mM ammonium formate pH 9.5 in water), 0.22µm cellulose acetate filtered (Costar Spin-X, Corning Inc., Salt Lake City, UT) and loaded onto an XBridge C18 column (3.5 µm, 2.1x 150 mm) from Waters, (Milford, MA) with 20 mM ammonium formate (NH4FA), pH 10 as buffer A and 85% ACN/15% 20 mM NH4FA as buffer B. The liquid chromatography was performed using a linear gradient at a flow rate 200 µL/min at 30 °C and the collection plate at 5 °C. Fractions were collected at 1-minute intervals and pooled into a total of 10 fractions based on the UV absorbance at 214 nm. Each of the 10 fractions was dried, redried from 200µl water (Optima LC-MS, Fisher Scientific, Fair Lawn, NJ) and reconstituted in 100 µL of 2% ACN/0.5% FA for nanoLC-MS/MS analysis.

### Enrichment of phosphopeptides

TiO_2_ enrichment was conducted using a TiO_2_ Mag Sepharose kit (from GE Healthcare). One aliquot of TMT 16-plex tagged peptides (800 μg) was reconstituted in 400 µL of binding buffer (1 M glycolic acid in 80% acetonitrile, 5% TFA). The TiO_2_ slurry (250 µL) was used and incubated with the sample for 30 minutes at 1,800 rpm on vortex mixer. After washing the beads with washing buffer (80% acetonitrile, 1%TFA), the phosphopeptides were eluted with 200 µL of elution buffer (5% ammonium hydroxide) twice. The eluted fractions were pooled, filtered with 0.22 µm spin filters (Costar), dried and reconstituted in 50 µL of 0.5% formic acid (FA) for subsequent nano scale LC-MS/MS analysis. In parallel, Fe-NTA enrichment was conducted using High Select Fe-NTA Phosphopeptide enrichment kit (Thermo). One aliquot of TMT 16-plex tagged tryptic peptides (800 μg) was reconstituted in the kit-provided binding/washing buffer (80% Acetonitrile, 1% TFA). The reconstituted peptides were then applied to spin columns containing equilibrated Fe-NTA beads and incubated at room temperature for 30 min. After washing the beads with binding/washing buffer, the phosphopeptides were eluted with 100 µL of the kit provided elution buffer (5% ammonium hydroxide) twice. The eluted fractions were pooled, filtered with 0.22 µm spin filters (Costar), dried and reconstituted in 50 µL of 0.5% formic acid (FA) for subsequent nano scale LC-MS/MS analysis.

### Nano-scale reverse phase chromatography and tandem MS (nanoLC-MS/MS)

Nano LC-MS/MS analysis was carried out using an Orbitrap Eclipse (Thermo-Fisher Scientific, San Jose, CA) mass spectrometer equipped with a nanospray Flex Ion Source using real-time search (RTS) SPS MS3 method coupled with the UltiMate3000 RSLCnano (Dionex, Sunnyvale, CA). Each reconstituted fraction (5 µL for global proteomics fractions) was injected onto a PepMap C-18 RP nano Viper trap column (5 µm, 100 µm’ 20 mm, Dionex) at 20 µL/min flow rate for rapid sample loading, and separated on a PepMap C-18 RP nano column (2 µm, 75 µm x 25 cm) prior to elution. In RTS mode, the *Drosophila melanogaster* Uniprot protein database with trypsin as the enzyme for real-time spectral database search. Dynamic exclusion parameters were set with ±10 ppm exclusion mass window. All data were acquired under Xcalibur 4.3 operation software in Orbitrap Eclipse (Thermo Fisher Scientific). For phosphoproteomics fractions, the Orbitrap Eclipse was operated in either multiple-stage activation (MSA)-based Synchronous Precursor Selection (SPS) MS3 method or neutral loss (NL) triggered MS3 method

### Data processing, protein identification and data analysis

All raw MS spectra were processed and searched using Proteome Discoverer 2.5 (PD2.5, Thermo). The *Drosophila melanogaster* database containing 41,409 entries was downloaded from Uniprot and used for database searches. Identified peptides were further filtered for maximum 1% FDR using the Percolator algorithm in PD 2.4 along with additional peptide confidence set to high and peptide mass accuracy ≤5 ppm. The TMT16-plex quantification method within Proteome Discoverer 2.5 software was used to calculate the reporter ion abundances in MS3 spectra that were corrected for the isotopic impurities. Both unique and razor peptides were used for quantitation. To confidently localize phosphorylated Ser, Thr, and Tyr residues, phosphoRS 3.0 mode was used.

### Bioinformatic and statistical analysis

TMT-normalized abundance values for both Fe-NTA and TiO2 phospho-peptides identified by neutral loss trigger and multistage activation were merged into one dataset. Phospho-peptide and total protein data were normalized to total peptide intensity for each respective dataset. To control for overall protein abundance changes (i.e., synthesis and degradation), phospho-peptide abundances were divided by parent protein abundance to generate normalized phosphopeptides values. Zero values were assigned to 0.01 prior to scaling to integer values and median normalization. To cluster phospho-sites detected in multiple peptides, abundance values of identical peptides with the same detected variable modifications were averaged. To determine differentially-regulated phospho-sites, Empirical Bayes moderated t-tests (Bioconductor LIMMA package in R Studio) were applied in a pairwise manner between conditions. Statistical significance was defined as Log2FC of >+/-0.5 and adjusted p-value of <0.05. To characterize phosphopeptides (i.e., Classes I-VI), phosphopeptides were compared for each condition and separated based on significant patterns of differential abundance. Principal component analyses and simple linear regressions were performed using R Studio.

Unbiased phospho-motif enrichment analysis was performed by collecting the 6 amino acids flanking either side of single phospho-modification from the *Drosophila melanogaster* Uniprot protein database. Our script calculates the frequency with which an amino acid appears in each position and determines an enrichment relative to a background dataset (in this case, the unchanging phospho-sites from our dataset). Heatmap plots were generated using R Studio (Fig. S4).

The above workflow was applied to phosphoproteomic datasets of Zhang et al. (2019; ‘CaNB2’ and ‘CnAct’) to determine phospho-S/T-P sites that are regulated by Calcineurin. To identify Calcineurin-regulated sites, differential abundance analysis was performed on Egg/Oocyte ratios from either control or Calcineurin-manipulated conditions. To identify statistically significant differences in proteins assessed by Western blots or development in ex vivo culture assays, students t-tests, Sidak’s/Dunnetts multiple comparisons test, or Mann Whitney tests were performed in GraphPad Prism 10.5.0.

### Fly stocks

Flies were reared on standard cornmeal/yeast/agar food and kept on a 12 hour light/dark cycle. Flies used in this study were as follows: Control flies used for phospho-mass spectrometry analysis were BL36304 control attP40 insertion flies crossed to BL57871. To achieve RNAi or overexpression of genes, females from the following lines were crossed to Maternal-alpha-Tubulin Gal4 (BL7063) males: *CDK1* RNAi (BL35350), *MAPK* RNAi (#1: BL36058; #2: BL34855), *CaNB2* RNAi (BL38971), UASp-Wee1-VFP (BL65390), UASp-CA-Calcineurin (‘CnAct’, Aigaki Lab, Zhang et al. 2019) and reared at 29°C. Additional lines used were *rl^1^* (BL386) and Nup358-GFP (Hampoelz et al. 2019). To generate spermless males for unfertilized egg collection, Control virgin females were mated to *sons of tudor* males (BL1786). To generate fertilized embryos, Control virgin females were mated to Canton S males.

### In vitro egg maturation (IVEM) assay

IVEM assay was adapted from Spracklen and Tootle (2013). 3-7 day old flies were fed dry yeast for 24 hours at 29°C (RNAi flies) or room temperature (*rl^1^*). Ovaries were dissected directly into room temperature IVEM medium (Schneider’s medium (Sigma) + 10% heat inactivated FBS (Gibco) + 1% penicillin streptomycin (Sigma)). Stage 12 egg chambers were manually freed from ovaries using pin vices (Ted Pella). Dissection time was limited to 30 minutes, and oocytes were inspected prior to beginning the assay; any oocytes that advanced beyond Stage 12 or lysed during the dissection period were discarded. Isolated Stage 12 egg chambers were deposited into a sterile 35mm culture dish with 1.5mL fresh IVEM medium warmed to room temperature, then grown in the dark at 25°C for 4 hours. Following IVEM, egg chambers were assessed morphologically for dumped nurse cells and dorsal appendage growth. Egg chambers were imaged on an ECHO Revolve fluorescence microscope under bright field settings (or FITC filter as necessary, Fig. S7F).

### Imaging

Oocytes of the indicated stages were collected from 3-7 day old flies fed on yeast for 24-48 hours at room temperature. Ovaries were dissected into PBS and then fixed in 4% paraformaldehyde in PBS for 12 minutes. Embryos were collected in 30-minute intervals, dechorionated with 50% bleach for 2 minutes, washed with egg wash buffer (0.4% NaCl, 0.03% Triton), and fixed in 4% formaldehyde and 50% heptane in PBS for 20 minutes. Following fixation, embryos were devitellinized: PBS and formaldehyde were removed and MeOH was added 1:1 with remaining heptane, and embryos were shaken vigorously. Media was removed and embryos were washed 3 times with 4mL MeOH. Oocytes and embryos were washed in PBS-T (0.1% Triton) and then stained with 20mM Hoechst 33342 at 1:1000 for 20 minutes at room temperature followed by three 15-minute washes in PBS-T. Oocytes and embryos were mounted in AquaPolymount and sealed with nail polish prior to imaging on a Zeiss LSM 880 inverted Axio Observer.Z1 microscope fitted with a Plan-Apochromat 40X/1.4 Oil objective. GFP was detected with a 488 laser and Hoechst was detected with a 405 laser.

### Gene Ontology, ortholog and disordered region predictions

Gene Ontology ‘Biological Process’ searches were conducted using Panther^[40]^. DIOPT^[41]^ was used to determine mouse and human orthologs of *Drosophila* phospho-regulated proteins and compare phospho-regulated proteins during fly and mouse^[38]^ oocyte maturation. PUNCH^[42]^ was used to predict intrinsically-disordered regions in Nup358 and ParSe 2.0^[43]^ was used to predict a residue’s likelihood to drive phase separation.

### Western blots and antibodies

Oocytes and embryos were collected as described above for proteomics methods, with the exception that oocytes were collected in PBS. Flash frozen oocytes were boiled directly in Laemmli buffer + BME (1uL per 1 egg chamber) for 5 minutes; embryos were briefly ground with pestles prior to boiling. Samples were centrifuged for 1 minute at max speed prior to loading on BioRad precast gels (10%, Fig. 5E or 4-20% gradient gels, all others). Gels were run in standard SDS-Tris-Glycine running buffer and transferred in Towbin + SDS for 80V for 30 minutes (Cyclin B, P-MAPK, Tubulin, CDK1, Gnu, Smaug) or Towbin lacking SDS for 2 hours at 200mAh (phospho-TP, phospho-SP). In Fig. 5E, western blots were performed on PFA-fixed samples. Oocytes were boiled for 20 minutes to remove crosslinks, and bands run at the same sizes for fixed vs. fresh samples (Cyclin B, CDK1, and Tubulin). Proteins were transferred to PVDF membranes, blocked for 30 minutes in Li-Cor Intercept buffer, and incubated with primary antibodies overnight at 4°C with gentle shaking. Membranes were washed with TBS-T (0.1% Tween) twice for 15 minutes at room temperature. Membranes were then treated with fluorescent secondary antibodies (Li-Cor: goat anti-rabbit IRDye 800 or 680, goat anti-mouse IRDye 680, or goat anti-guinea pig IRDye 800) for 1 hour at room temperature in the dark. Membranes were washed twice with TBS-T and then washed once with TBS. Blots were then imaged on either a Li-Cor Odyssey CLx (Fig. 5E) or Li-Cor Odyssey M (all others) prior to quantification of fluorescence intensity in ImageStudioLite Version 5.2. If applicable, post-imaged membranes were incubated in Coomassie Brilliant Blue (CBB) until membranes were blue and then destained in 50% MeOH, 1% acetic acid until bands were visible.

Antibodies were used at the following concentrations: rabbit anti-CDK1 (Sigma, ‘PSTAIR’, 1:1000), rabbit anti-phospho-MAPK-T302/Y304 (Cell Signaling Technologies, 1:5000), mouse anti-alpha Tubulin (Sigma, 1:5000), rabbit anti-alpha Tubulin (Cell Signaling Technologies, 1:5000), guinea pig anti-Smaug (Howard Lipshitz, 1:10000), guinea pig anti-Gnu (Masatoshi Hara and Terry Orr-Weaver, 1:10000), mouse anti-Cyclin B (Developmental Studies Hybridoma Bank, DSHB, University of Iowa, ‘F2F4’, 1:50), mouse anti-phospho-Threonine-Proline (Cell Signaling Technologies, 1:1000), rabbit anti-phospho-Serine-Proline (Cell Signaling Technologies, 1:1000). All secondary antibodies were used at 1:10000.

## Results and Discussion

### The phosphoproteome changes dynamically during oocyte maturation and egg activation, but is not impacted by fertilization

We sought to understand the developmental regulation of phosphorylation state amongst maternally supplied proteins. We performed quantitative phosphoproteomic characterization of Stage 12 and 14 oocytes and on newly fertilized embryos or activated, unfertilized eggs (<30 minutes after egg laying) (Fig. 1A). Phosphopeptide abundances were normalized to parent protein abundances to control for changes in overall protein levels (see methods for full normalization pipeline). PCA analysis confirmed that Stage 12, Stage 14, and Activated egg samples formed distinct clusters (Fig. 1B). Pairwise linear regression consistently yielded R^2^ values of 0.88 and higher between replicates, indicating highly reproducible results (Fig. S1A-D).

**Figure 1:**
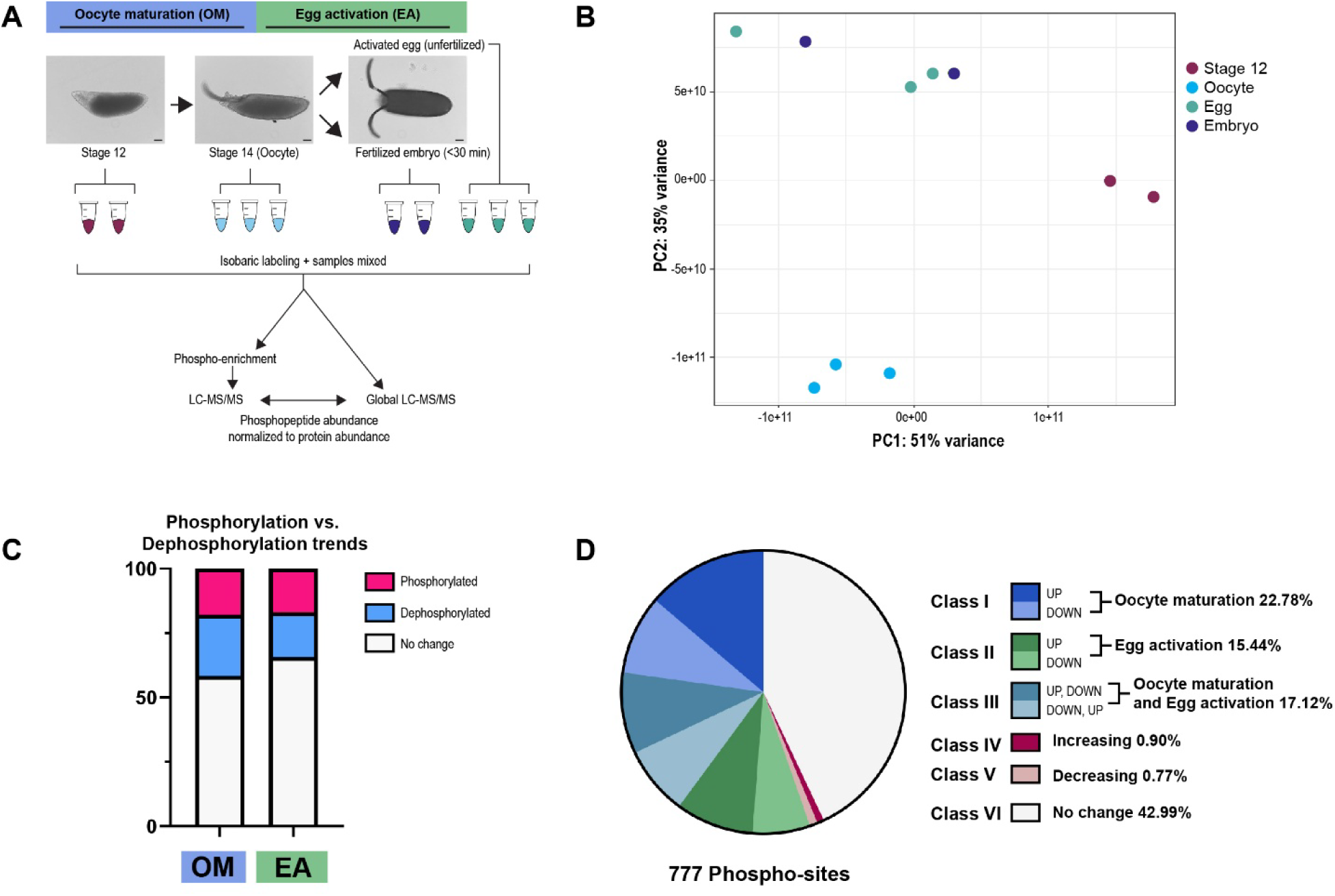
Phosphoproteomic mass spectrometry analysis of oocyte maturation (OM), egg activation (EA), and fertilization. A) Experimental overview of samples collected for multiplexed phospho-LC-MS/MS analysis, for which approximately 500 oocytes or eggs were used per replicate. Scale bars = 50μm. B) Principal component analysis of normalized phosphopeptides. C) Phosphorylation versus dephosphorylation observed during OM and EA. D) Proportion of phosphopeptides that are differentially regulated during OM, EA, or both.

After normalization and clustering of phosphopeptides, 777 phosphopeptides corresponding to 356 proteins were subjected to differential analysis. We also analyzed the global proteome changes and found similar trends to former analyses^[44]^ (Fig. S2 and Table S1, comparison to Kronja et al. 2014). Phospho-normalized values were median normalized and analyzed by empirical Bayes moderated t-tests using LIMMA^[5, 44]^. In total, 41.6% of phosphopeptides were significantly changed during OM (i.e. Stage 14-12 comparison), with a slight bias towards dephosphorylation of substrates (Fig. 1C). During EA (i.e. Egg-Stage 14 comparison) 34.2% of phosphopeptides were differentially phosphorylated, with phosphorylation and dephosphorylation occurring at comparable levels.

We next grouped phospho-proteins into classes based on when they showed differential changes or lack thereof (Fig. 1D). Class I phospho-sites (22.78%) were significantly changed during OM, but unchanged during EA. Class II sites (15.44%) were changed during EA, but showed no changes during OM. Class III sites (17.12%) were changed during both OM and EA, in opposing directions (e.g. phosphorylated in OM, dephosphorylated in EA, or vice versa). Class IV sites (0.90%) were phosphorylated during both OM and EA. Class V sites (0.77%) were dephosphorylated during both OM and EA. Finally, Class VI sites (42.99%) were unchanged across OM and EA.

Shortly after *Drosophila* EA is initiated by mechanical pressure during transit of the egg through the oviduct^[28]^, the egg is fertilized^[45]^. In mammals, sperm soluble factors stimulate EA^[46]^. While sperm does not provide the initial trigger for EA in *Drosophila*, we hypothesized that a soluble factor (e.g., kinase, activator, inhibitor) could contribute to phospho-signaling at EA by being released during degradation of the sperm tail’s plasma membrane and mitochondrial derivative^[47, 48]^. We thus compared the phosphoproteomes of activated eggs that were either fertilized or unfertilized. Surprisingly, we found no evidence of significantly different phosphopeptide abundances between eggs and embryos (Fig. S3A); this result is reflected in our PCA analysis, as egg and embryo clusters overlap (Fig. 1B). Relaxing the significance threshold (e.g. 0.1 adjusted p value) revealed only two changed phospho-proteins between eggs and embryos: heterochromatin protein 1 (HP1/Su(var)205) and CanA-14F, the catalytic subunit of Calcineurin. At the proteome level, we found only one significant change to MetRS (Fig. S2C).

To corroborate our finding that activated and fertilized eggs have equivalent phosphorylation profiles, we analyzed the degree of phosphorylation changes during EA in both conditions by comparing the log2 fold changes of phosphopeptides between Stage 14 oocytes versus embryos or eggs. The phosphorylation changes at EA in unfertilized eggs were nearly identical to those of fertilized eggs. We analyzed the phospho-proteins with log2 fold changes greater than 0.5 in an attempt to find sperm contributions, but none were significant (Fig. S3B).

We acknowledge that our stringent identification and quantification criteria may have eliminated relevant targets, or that our analysis may lack the sensitivity necessary to detect changes to particularly low-abundance phospho-proteins. However, we emphasize that the same analysis found abundant changes between the other sample types in the same multiplexed experiment. We therefore conclude that sperm does not induce significant changes to the phosphoproteome in the early embryo and that phospho-changes during EA are predominantly driven by maternal factors. This suggests the interesting possibility that in vertebrate systems sperm serves to simply trigger EA, but does not impact cellular signaling until the zygotic genome activates. We suspect that the maternal ooplasm is the driving developmental force and will act upon sperm if present, but development can proceed unimpeded until a developmental block is imposed by the need for the sperm’s centriole during the first mitotic division^[49]^ or the sperm’s genome contribution is needed.

### CDK1/MAPK substrates are dephosphorylated during egg activation

The phosphoproteome is remodeled during EA, and phosphorylation changes are dependent on Calcineurin^[5]^. However, the kinases that establish a mature oocyte’s phosphorylation profile were unknown. To identify these kinases, we assessed phospho-site motifs for predicted kinases. We first determined that our analysis detected phospho-changes during EA as expected and that phospho-regulated proteins belong to expected biological processes (Fig. 2A-B). We then analyzed differentially-abundant phosphopeptides using an unbiased phospho-motif enrichment analysis (Fig. S4). Phospho-S/T-P is the preferred target of MAPK/Erk and CDK1/cdc2^[50]^. In *Drosophila*, Erk, p38, and JNK MAPK deactivate during EA^[51]^. For the current analysis, we focused on Erk and refer to it as ‘MAPK’. At the same timepoint as MAPK’s decrease in activity, CDK1’s activity increases^[5]^, possibly reflecting a strong need for CDK1 activity for polar body formation/degradation and initiation of embryonic mitoses^[52]^.

**Figure 2:**
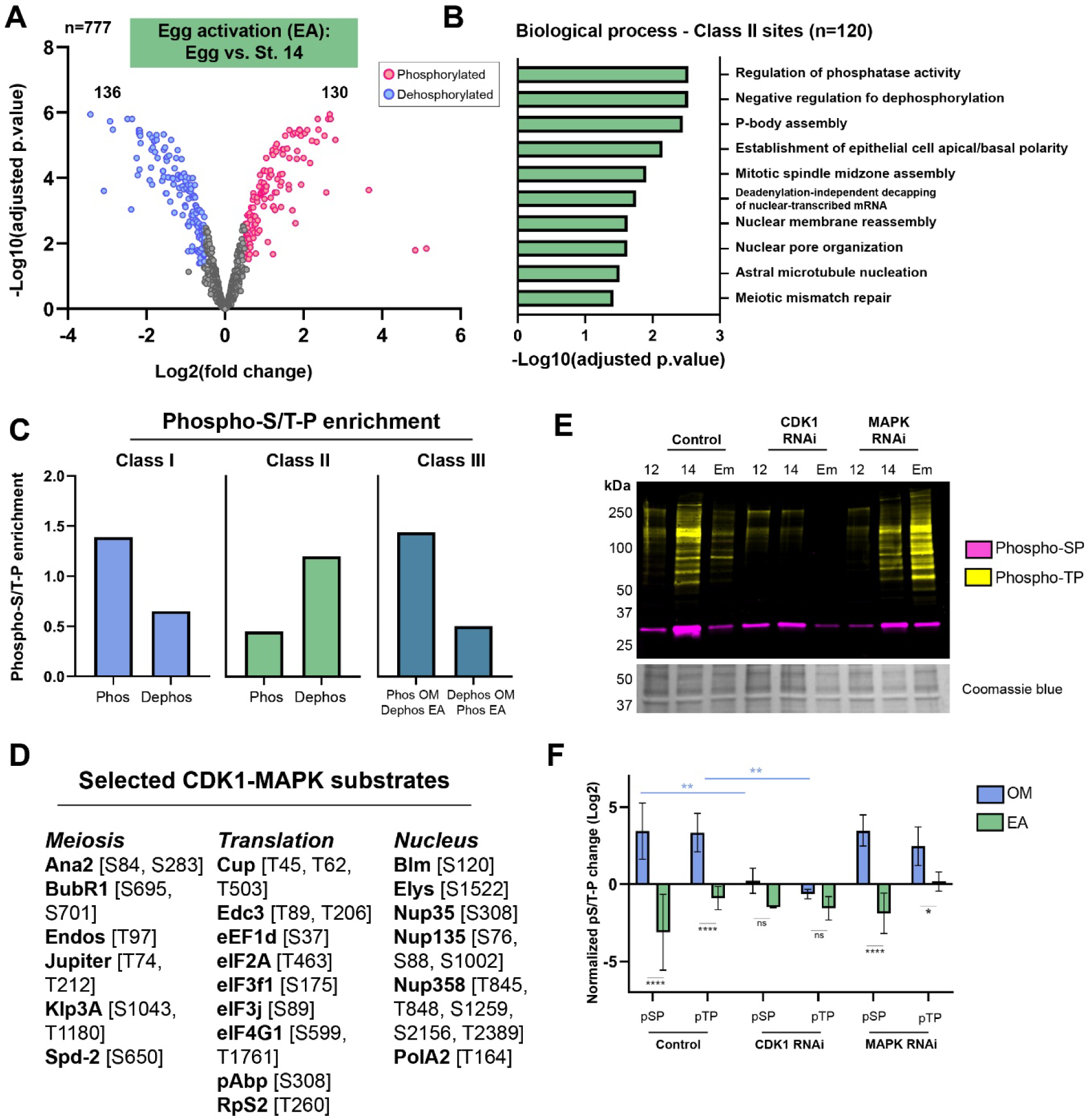
CDK1-MAPK substrates are dephosphorylated during egg activation. A) Volcano plot of phosphopeptide changes during EA. 266/777 phosphopeptides from 163 proteins are differentially regulated, with 120 changes specific to EA. B) GO analysis performed on proteins whose phospho-sites change only during EA; highly significant terms for Biological Process are displayed; for full GO output, see File S2. C) Enrichment of phospho-S/T-P motifs among differentially regulated phosphopeptides for Class I, II, and III substrates compared to the unchanging pool of phospho-sites. Phospho-S/T-P sites are enriched in sites that are phosphorylated during OM (‘Phos’, Class I, III) and dephosphorylated during EA (‘Dephos’, Class II, III). E) Selected phospho-S/T-P substrates that change significantly during OM, EA, or both as related to Meiosis, Translation, and Nuclear processes. E-F) Western blot analysis of phospho-S/T-P proteins (Phospho-SP, magenta, Phospho-TP, yellow) during OM and EA for control, CDK1 RNAi, and MAPK RNAi conditions. Phospho-S/T-P levels were normalized to Coomassie blue intensity and were found to increase during OM and decrease during EA in control and MAPK RNAi conditions. CDK1 RNAi oocytes fail to acquire their Stage 14 phospho-signatures during OM and Phospho-S/T-P levels are severely reduced in embryos. All samples were compared by Sidak’s multiple comparison test; significance or lack thereof for relevant conditions are displayed. The experiment was repeated three times. *p<0.05, **p<0.01, ****p<000.1, ns=p>0.05.

Phospho-S/T-P substrates were highly represented across all conditions analyzed (Fig. 2C). CDK1-MAPK substrates make up 92% of phosphorylated substrates (Class I) during OM (Fig. S4A). Complementary to this observation, 79% of sites dephosphorylated during EA (Class II) bear S/T-P motifs (Fig. S4B). Within Class III phosphorylation sites, 95% of those that are phosphorylated during OM and dephosphorylated during EA are represented by CDK1-MAPK substrates (Fig. S4C). Significantly-changed CDK1-MAPK substrates were related to meiotic progression, translational reprogramming and RNA handling, and nuclear processes, among others (Fig. 2D). Together, these results suggest that CDK1 and/or MAPK actively phosphorylate some substrates and maintain phosphorylation on others during OM, and a portion of these phosphorylation events are counteracted during EA.

As an orthogonal method to assess the roles of MAPK and CDK1 in remodeling the phosphorylation landscape, we utilized antibodies that are specific to phospho-TP or phospho-SP motifs (Fig. 2E). In control samples, we found dramatic increases in phospho-S/T-P phospho-proteins during OM, and a strong decrease in intensity during EA (Fig. 2E-F). We utilized germline specific knockdown of MAPK and CDK1 (Fig. S7A,C) to test the genetic requirements for these kinases in regulating S/T-P substates during OM and EA. Stage 12 oocytes depleted for CDK1 have phospho-profiles comparable to controls, but Stage 14 oocytes failed to acquire phospho-S/T-P marks and instead remain largely unchanged from the Stage 12-state (Fig. 2E-F). In activated fertilized eggs, all detectable phospho-TP motifs were missing, despite unchanged total protein abundance. Thus, CDK1 is a major regulator of phosphorylation signatures during OM and EA. Interestingly, when MAPK is knocked down, phospho-profiles look similar to control samples, and somewhat elevated in early embryos, suggesting either a minor role for MAPK or compensation by another kinase.

### Phosphorylation of CDK1-MAPK substrates during oocyte maturation is counteracted by Calcineurin during egg activation

CDK1 is classically studied in the context of metaphase arrest, whereby dephosphorylation of its substrates Greatwall and Endos permits the activation of PP2A and stimulation of APC/C.^[53, 54]^ During mouse EA, CDK1 deactivation is thought to be calcium dependent^[55]^; we therefore tested whether CDK1 substrate dephosphorylation requires the relevant calcium-activated phosphatase Calcineurin. We assessed CDK1-MAPK substrates (via phospho-S/T-P western blotting) in early embryos lacking Calcineurin (Calcineurin RNAi) or expressing constitutively-active Calcineurin (CA-Calcineurin^[14]^) (Fig. 3A-B). Calcineurin depletion caused significant increases in both phospho-SP and phospho-TP substrate abundance (Fig. 3A-B), indicating their retention on target proteins during EA. In early embryos expressing CA-Calcineurin, we found a trending, but nonsignificant, decrease in phospho-SP (p=0.1); phospho-TP substrates decreased slightly (p=0.7). Thus, Calcineurin is necessary, but not sufficient, for dephosphorylation of CDK1/MAPK substrates.

**Figure 3:**
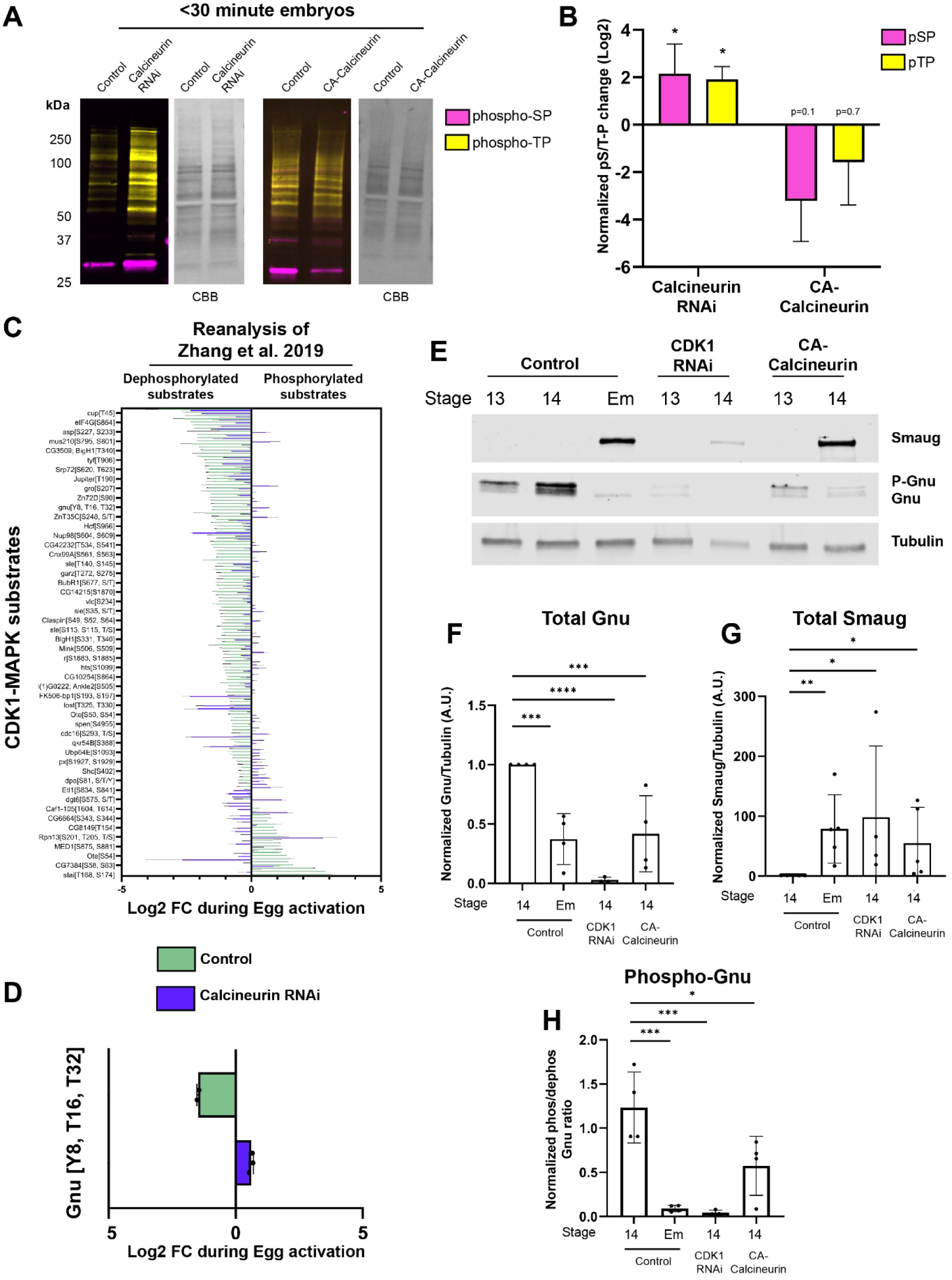
Calcineurin counteracts phosphorylation of S/T-P sites during EA and promotes embryonic translation via Gnu. A-B) Western blots of embryos 30 minutes-post egg laying from control, Calcineurin RNAi and CA-Calcineurin flies. Phospho-SP (magenta) and phospho-TP (yellow) signal was normalized to Coomassie brilliant blue (CBB) intensity and plotted as the ratio of the indicated condition compared to control. Levels were compared to controls via Mann Whitney test, *p<0.05. The experiment was performed three times. C) Calcineurin-regulated phospho-S/T-P sites identified via reanalysis of Zhang et al. 2019. Green bars indicate the Log2 fold change of substrates during control EA, while indigo bars indicate the same site in Calcineurin RNAi samples. All changes shown have statistically-significant differences in Log2 fold changes between control and Calcineurin RNAi. D) Dephosphorylation of Gnu’s T16 and T32 CDK1 sites is Calcineurin dependent; values are lifted from panel C. E-H) Western blots and quantification for Smaug, Gnu, and Tubulin in oocytes and embryos. In control oocytes (Stage 14), Smaug is never detected, but is present in CDK1 RNAi oocytes and oocytes overexpressing CA-Calcineurin. Gnu is primarily phosphorylated in control oocytes and is dephosphorylated and degraded in embryos. Gnu is prematurely dephosphorylated and degraded in CDK1 RNAi and CA-Calcineurin oocytes. For overexposed blots that display dephosphorylated Gnu in CDK1 RNAi Stage 14 oocytes, see Fig. S8H. The experiment was performed four times per condition and analyzed via Dunnetts multiple comparisons test (Phospho-Gnu and total Gnu, F and H) and Dunns multiple comparisons test (Smaug, G). *p<0.05, **p<0.01, ***p<0.001, ****p<0.0001.

Our former proteomic analysis of EA revealed that numerous substrates are regulated by Calcineurin^[5]^, including those whose dephosphorylation does not normally occur in Calcineurin RNAi conditions or is exaggerated in CA-Calcineurin conditions. We thus reanalyzed our former dataset to assess whether MAPK/CDK1 targets are among Calcineurin-regulated substrates. We applied our stringent phospho-normalization pipeline to both Calcineurin RNAi and CA-Calcineurin datasets and found a surprisingly mild effect on phospho-proteins when CA-Calcineurin is overexpressed (Fig. S5A-B; a pool of 80 phospho-sites that trend toward significance at p<0.2 and 28 affected phospho-S/T-P sites). Therefore, CA-Calcineurin overexpression is insufficient for widespread changes to maternal phospho-proteins during EA, possibly because overexpressed CA-Calcineurin offers little additional phosphatase activity to the surge of endogenous Calcineurin. In contrast, we found a strong requirement for Calcineurin to dephosphorylate CDK1-MAPK substrates. Of 1726 normalized phospho-sites, we identified 554 (32.1%) changes in control conditions. Of those 554 sites, 312 (56.3%) phospho-sites are significantly altered in Calcineurin RNAi activated eggs. We assessed the pool of misregulated phospho-sites for the presence of phospho-S/T-P motifs. Of the 312 misregulated sites, 149 (47.8%) sites were CDK1-MAPK sites, belonging to 100 total proteins (Fig. 3C). Among these phospho-sites, 79.2% behaved like Calcineurin substrates: these sites were dephosphorylated in control conditions, but either stayed constant or increased in their phosphorylation in Calcineurin RNAi conditions. A smaller class of phospho-sites (20.8%) increased normally in controls, but were further dephosphorylated in Calcineurin-depleted eggs.

In *Drosophila* and *Xenopus*, CDK1 substrate Fzy/Cdc20 is dephosphorylated via PP2A at EA^[33]^. Fzy is an activator of the APC/C^[56]^, and its role is tied to degradation of Cyclin B^[57]^. Calcineurin is necessary for normal phospho-regulation of Fzy’s CDK1-sites (Fig. 3C; Fzy T78 and T82), likely placing Calcineurin upstream or parallel to PP2A and APC/C activation in the regulatory hierarchy of CDK1 targets at EA. It will be interesting to delineate PP2A and Calcineurin’s distinct and overlapping targets. Calcineurin is required for fertility and release from meiotic arrest; we found that almost 50% of Calcineurin’s effectors are CDK1-MAPK targets, suggesting that failed modification of these targets via Calcineurin underlies this fertility defect.

### CDK1 and Calcineurin antagonize embryonic translation initiation via Gnu

Both meiotic and translation regulators are phospho-regulated by CDK1 and Calcineurin during EA (Fig. 2D and 3C). Mature oocytes lacking CDK1 fail to establish metaphase arrest: they remain locked in a prophase-like state and retain the GV beyond Stage 12^[31]^. CDK1’s effect on translation has not been studied in this context. We therefore assessed translation events at EA by western blotting for Smaug, one of the first translated proteins after EA (Fig. 3E). Smaug destabilizes and clears maternal mRNAs, which is essential for downstream activation of the zygotic genome^[58]^.

Smaug is never detected in Stage 13 or 14 oocytes from control flies, but is abundant in early embryos (<30 minutes after egg laying, Fig. 3E, G). In Stage 13 oocytes lacking CDK1, Smaug is absent as expected (Fig. 5E). However, Smaug is present in Stage 14 CDK1 RNAi oocytes. The same assay found increased Smaug protein in Stage 14 oocytes expressing CA-Calcineurin (Fig. 3E). We note that for both manipulations, the absolute levels of Smaug detected in oocytes varied (Fig. S8H for additional examples), possibly due to variability in knockdown or overexpression efficiency between experiments. However, we note the absence of Smaug in control Stage 14 oocytes and highlight that the Smaug detected in CDK1 and Calcineurin-manipulated oocytes indicates premature onset of the embryonic translation program.

Translation is initiated during *Drosophila* EA via PNG, whose subunit Gnu must be dephosphorylated to complex with its members and promote Smaug translation. Gnu contains putative CDK1 sites^[36]^, two of which (Phospho-T16 and T32) are dephosphorylated in a Calcineurin-dependent manner during EA (Fig. 5C-D and Zhang et al. 2019). To determine whether the CDK1-Calcineurin-dependent Smaug translation is mediated via PNG, we analyzed Gnu phosphorylation in CDK1 RNAi and CA-Calcineurin Stage 14 oocytes. We found that Gnu is dephosphorylated both when CDK1 activity is reduced and when Calcineurin activity is increased (Fig. 3E, H). Interestingly, total Gnu levels are reduced in both CDK1 RNAi and CA-Calcineurin oocytes (Fig. 3E-F). Gnu levels normally drop within the first 1-2 hours of embryogenesis^[59]^ and the observed reduction of total Gnu in our CDK1 and Calcineurin-manipulated oocytes therefore suggests that embryonic degradation of Gnu occurs prematurely.

Our results demonstrate an in vivo requirement for CDK1 to restrict premature dephosphorylation and degradation of Gnu. Loss of Gnu’s phosphorylation by CDK1 promotes downstream PNG-mediated Smaug translation in the oocyte (Fig. 3E), and oocyte Gnu levels fall to a similar level as in activated embryos when CDK1 or Calcineurin activity is modified. Calcineurin is likely the phosphatase that dephosphorylates Gnu: Gnu dephosphorylation is dependent on Calcineurin^[5]^, and boosted Calcineurin activity promotes its dephosphorylation and degradation, leading to Smaug translation in the Stage 14 oocyte. In addition to Gnu, our analysis identifies several translation regulators which may be required to restrict or promote specific translational events during EA: CDK1-MAPK sites on initiation factors eIF3f1 and eIF4G1, as well as ribosomal subunit RpS2, are phosphorylated during OM (Fig. 2D). RNA handling proteins, like Cup and Edc3, are dephosphorylated during EA (Fig. 2D); among these, eIF4G1 and Cup dephosphorylation requires Calcineurin (Fig. 3C). Our analysis highlights novel targets of CDK1-MAPK that influence EA progression and further reveals that misregulation of translational events by CDK1 or Calcineurin can mistime EA.

Together, our analysis finds that cellular events that are parallel to meiotic processes can progress beyond release from prophase arrest. Thus, the same regulator (i.e., CDK1) that is needed to release prophase arrest is needed to restrict translational events that normally occur following establishment and release from metaphase arrest. In other words, translational activation at EA progresses as if metaphase arrest was relieved because this arrest had never been established. We propose that meiotic and translation processes, despite having a shared upstream regulator, present different developmental trajectories if CDK1 fails to deploy.

### Phospho-proteins are primed for egg activation during oocyte maturation

The events of EA rely on the accumulation and regulation of macromolecules during earlier oogenesis. Most maternally-provided proteins are synthesized prior to their deposition in the oocyte; the Stage 12 oocyte is the first stage after which deposition has completed. We thus compared phospho-protein changes prior to EA beginning at Stage 12.

### Substrates regulated during both OM and EA

By coupling the analysis of OM and EA, we could identify phosphopeptides that are differentially regulated during either transition or both. Almost 20% of phosphopeptides (146 on 109 proteins) are regulated across both transitions: a small subclass of phosphopeptides stably increase (Class IV) or decrease (Class V) during both transitions. Interestingly, the majority of phosphopeptides regulated during both transitions have opposing abundance changes across developmental time (Class III) (Fig.1D, Fig. 4A). Class III sites represent proteins that are phosphorylated during OM and then dephosphorylated during EA, or vice versa. GO analysis revealed that Class III substrates are largely related to handling of stored mRNA or oocyte patterning (Fig. 4B). For both OM-specific and OM-EA regulated phospho-proteins, CDK1/MAPK motifs were highly represented (Fig. 2C).

**Figure 4:**
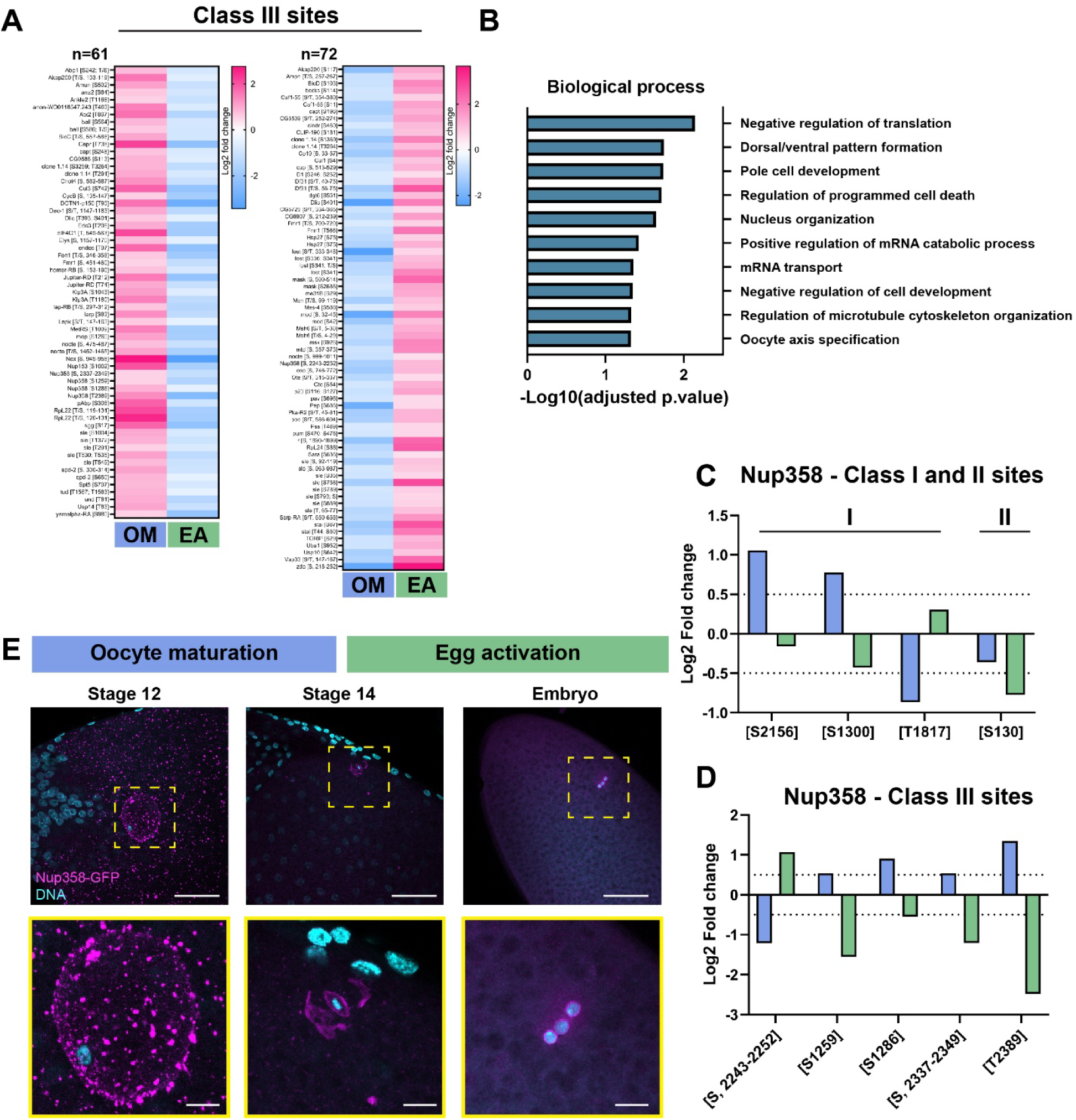
Class III substrates, including Nup358, are regulated during both OM and EA. A) Heatmaps of Class III substrates, which undergo differential regulation in opposing directions between OM and EA. B) GO analysis performed on shared substrates; highly significant terms for Biological Process are displayed (full GO output can be found in File S2. C-D) Nup358 bears Class I, II, and III sites. E) Maximum intensity projections of confocal sections of fixed Stage 12 and 14 oocytes and embryos stained for DNA (Hoechst, cyan) and expressing endogenously-tagged Nup358-GFP (magenta). Nup358 is abundant in the Stage 12 ooplasm, with enrichment on the germinal vesicle. In Stage 14, Nup358 is concentrated around the oocyte nucleus in a pattern that is consistent with close apposition to the metaphase I spindle. In embryos, Nup358 is enriched on polar bodies. Images were brightened in Fiji to aid in visualization. Unadjusted images, and additional examples of each stage, are shown in Fig. S4F-H. For each condition, 2 (embryo) to 3 (Stage 12 and 14) independent slides were imaged and assessed. Scale bars = 50μm (top row), 10μm (bottom row).

A phospho-site that is regulated during one transition may belong to a parent protein with additional sites that are regulated during another transition (i.e. one protein may house both Class I and II phospho-sites, and/or Class III sites). Our analysis identified 27 such proteins (Table S2), including those bearing CDK1-MAPK motifs. One of these proteins, Nup358, undergoes phase separation during fly oogenesis to support the assembly of precursor nuclear pore complexes (annulate lamellae) in a Ran-dependent manner^[60]^, which are thought to provide pre-assembled nuclear pore complexes to meet the need for rapid nuclear synthesis during early embryonic mitoses. We hypothesized that Nup358’s phosphorylation state might affect its propensity to phase separate or localize to particular structures.

We mapped phospho-regulated sites to intrinsically disordered regions (IDRs) within Nup358’s Alphafold2 predicted structure ^[42, 43, 61]^ (Figure S5). Interestingly, half of Nup358’s phospho-regulated sites are predicted to occur in IDRs, including in CDK1-MAPK sites (T848 and S2156, Fig. S5B-E). Further prediction of a residue’s propensity to phase separate using ParSe revealed both S1300 and S2156 as prone to phase separation (Fig. S5C, E), for which S2156 is a CDK1-MAPK site.

To determine whether Nup358 localization correlates with its phospho-states, we imaged oocytes and embryos expressing GFP in-frame with endogenous Nup358^[60]^. During Stage 12, Nup358 is present in multiple pools: phase separated puncta are abundant in the ooplasm ^[60]^ and a second population of Nup358 is present on the surface of the GV^[62]^ (Fig. 4E). In Stage 14 oocytes, Nup358 shows reduced ooplasmic foci, but is enriched around the Metaphase I spindle (Fig. 4E). In the newly laid embryo (<30 minutes), cytoplasmic foci were further reduced, and Nup358 was enriched around polar bodies (Fig. 4E). While our analysis does not account for which phospho-forms of Nup358 localize to specific structures, our results reveal changes to its bulk and sub-cellular localization in a pattern that mirrors its phospho-state changes. We highlight Nup358 as an example of 27 proteins bearing both Class I and II sites (Table S2), whose phospho-regulatory trends warrant further investigation

Phosphorylation is known to dictate the phase-separation behavior of proteins with disordered regions^[63, 64]^, including in *Xenopus* embryos, where changes to proteins’ phase separation occur via CDK1-dependent phospho-changes of their IDRs^[65]^. It is unclear if phosphorylation dynamics drive protein localization during *Drosophila* OM and EA, and what affect localization changes may have on development. We speculate that disrupted phospho-regulation of Nup358 at its phase separation-promoting sites could underlie the failed release from Prophase I arrest in CDK1 RNAi oocytes if GV breakdown is a prerequisite for metaphase establishment.

### Substrates regulated specifically during OM

We identified 323 significantly-changed phosphopeptides belonging to 203 proteins between Stage 12 and Stage 14 (OM) (Fig. 5A). Of these, 185 are dephosphorylated and 138 are phosphorylated. 177 out of 323 phosphopeptides change uniquely during OM (Class I). GO enrichment analysis of Class I sites (Fig. 5B) for ‘Biological Process’ found terms related to the handling of stored maternal mRNA and meiotic progression.

**Figure 5:**
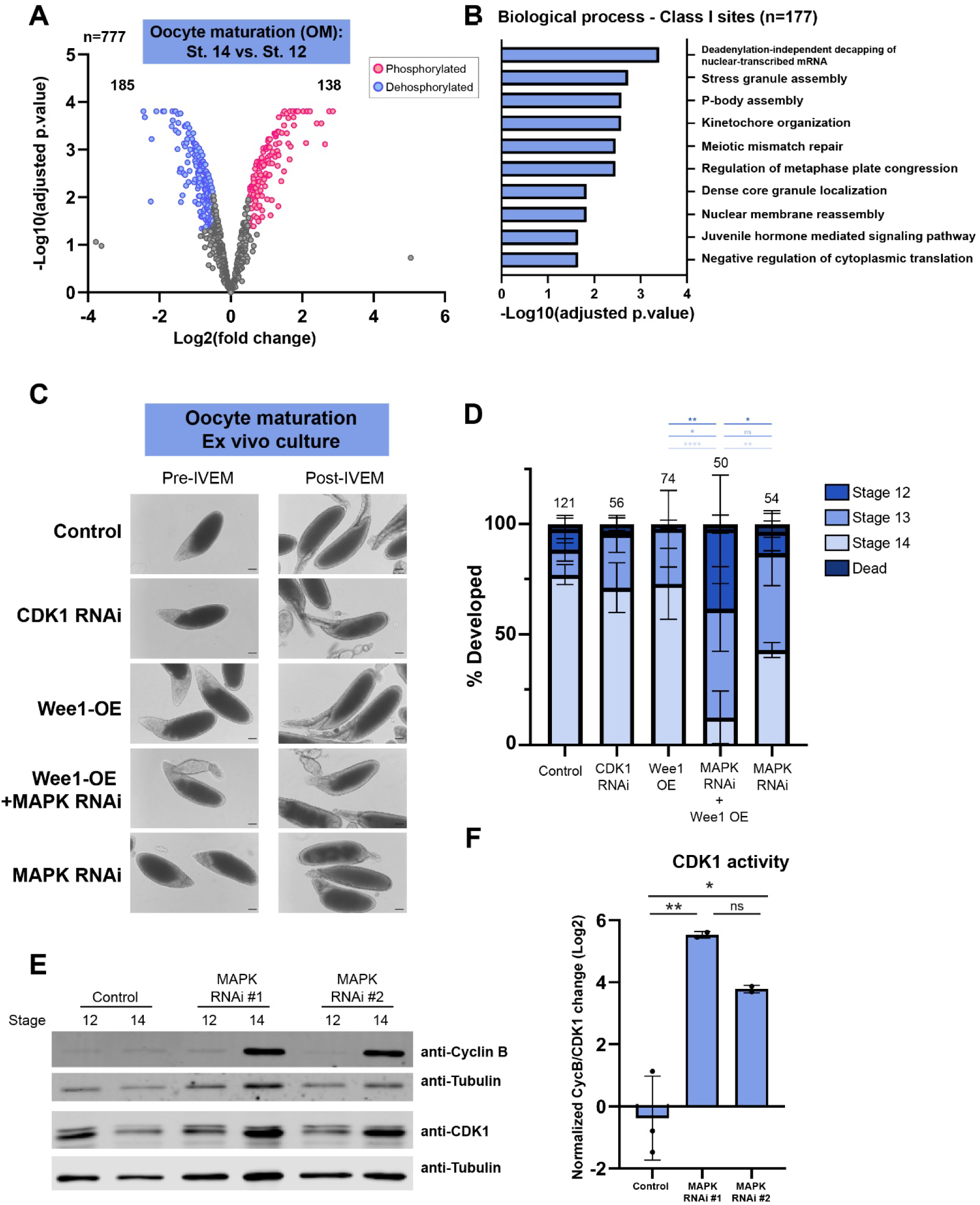
CDK1 activity rises in response to reduced MAPK signaling. A) Volcano plot of phosphopeptide changes during OM. 323/777 phospho-sites on 203 proteins are differentially regulated, with 177 changes specific to OM. B) GO analysis performed on proteins whose phospho-sites change only during OM; highly significant terms for Biological Process are displayed; for full GO output, see File S2. C) Representative bright field images of Stage 12 oocytes from the indicated conditions before and after in vitro egg maturation (IVEM). Scale bars = 50μm. For additional examples and validation of genetics tools, see Fig. S7. D) Quantification of IVEM assays. Differences in OM efficiency were analyzed via Tukey’s multiple comparisons test. The number above each bar indicates the number of oocytes used in the analysis, and the experiment was repeated at least three times per condition. E-F) Western blot and quantification for Cyclin B and CDK1 in Stage 12 and 14 oocytes from controls or two independent MAPK RNAi conditions. The normalized change of Cyclin B/CDK1 from Stage 12 to Stage 14 is plotted and differences were assessed via Tukey’s multiple comparisons test. The experiment was repeated twice. For all statistics, *p<0.05, **p<0.01, ***p<0.001, ****p<0.0001, ns=p>0.05.

Our results indicate that there are multiple pools of maternally-deposited proteins. This includes those whose phospho-signatures match the “arrest state” (Class II; these proteins are deposited in a phospho-form that is compatible with EA), another pool that rapidly undergoes phosphorylation changes during OM, but does not change during EA (Class I; these substrates are likely not used for EA, but are relevant for the processes required for OM), and finally, proteins that are placed in the Stage 12 oocyte in a phospho-state that matches the identity of the activated egg’s proteins, with a transient change in phospho-state before EA (Class III). These proteins may perform their functions during OM, but then turn off to maintain oocyte arrest, and reactivate upon EA (or vice versa). Our data suggests a paradigm in which oocytes recycle the same protein for different developmental processes in a phospho-regulated manner. Protein phospho-state may also underlie differential functions of proteins whose expression is not restricted to oocytes (e.g. pleiotropy). The utilization of phospho-regulation during EA, rather than transcriptional or translation changes, may reflect the need for a rapid mechanism to change a gene product’s function. Further, phosphorylation changes may be relatively energetically favorable for the dormant oocyte compared to transcription or translation changes.

### MAPK regulates oocyte morphogenesis and its loss is compensated for by CDK1

We next assessed the broad genetic requirements for OM by monitoring oocyte morphogenesis. MAPK RNAi ovaries contain a mix of both morphologically normal Stage 14 oocytes and those completely lacking dorsal appendages (Fig. S7D). To determine whether failed dorsal appendage formation occurs due to loss of MAPK during OM, we employed a modified in vitro egg maturation (IVEM) assay^[66]^ (Fig. 5C), whereby Stage 12 explants are cultured in the presence of FBS and antibiotics for 4 hours, during which they mature into Stage 14 oocytes. In control oocytes, morphologically normal Stage 14 oocytes develop 77.1% of the time (Fig. 5C-D). We assessed the need for MAPK by performing IVEM on egg chambers from our validated MAPK RNAi flies and those bearing a hypomorphic allele (*rl^1^*, Fig. S7G; validation of reduced MAPK, Fig. S7B). In MAPK RNAi conditions, Stage 12 egg chambers appear morphologically normal, but only 42.9% mature to Stage 14, while 43.9% mature to Stage 13, with failed or stunted dorsal appendage growth (Fig. 5C-D). In the hypomorphic *rl^1^* background, we observed a similar failure to mature into Stage 14 oocytes (Fig. S7G). Oocytes appeared smaller in size and had increased frequency of failed OM, as well as examples of abnormal dorsal appendages. We conclude that MAPK signaling in the egg chamber is needed for faithful oocyte morphogenesis and dorsal appendage growth.

We performed the same IVEM assay on oocytes with CDK1 knocked down. As an independent approach to deactivate CDK1, we overexpressed VFP-tagged Wee1 (Wee1-OE), a tyrosine kinase that negatively regulates CDK1 by phosphorylating its inhibitory Y15 site^[67]^ (Wee1-OE expression was verified by fluorescence imaging, Fig. S7F). We were surprised to find that, in both cases, Stage 12 oocytes grow into Stage 14 oocytes in culture over 70% of the time, with no statistical difference from control counterparts (Fig. 5D). Despite the failure to release Prophase I arrest^[31]^, oocytes appear grossly normal, but we observed a subpopulation of CDK1 RNAi and Wee1-OE oocytes with a clear region around the periphery, likely indicating reorganization of lipid and yolk (Fig. S7D).

We were puzzled by the inability of MAPK RNAi oocytes to fully mature when their phosphorylation profiles appeared similar to controls. We hypothesized that, because of their shared substate preference, CDK1 and MAPK could compensate for each other. To test this idea, we performed our IVEM assay on flies with both MAPK and CDK1 activity depleted. To circumvent oocyte lethality from double knockdown of CDK1 and MAPK, we overexpressed Wee1 (Wee1-OE) in MAPK RNAi flies. Compared to Wee1-OE or MAPK RNAi alone, Wee1-OE;MAPK RNAi showed a dramatic reduction in OM (Fig. 5D): only 12.38% of oocytes matured to Stage 14, and others were present in a Stage 12 or 13-like state. Thus, inhibition of CDK1 via Wee1-OE in combination with MAPK knockdown dramatically reduces the efficiency of OM.

The synergistic decrease in oocyte development when CDK1 and MAPK activity are both depleted suggested that CDK1 may compensate for reduced MAPK signaling. We thus assessed levels of CDK1 and its binding partner Cyclin B to assess Maturation Promoting Factor (MPF) activity. In Stage 12 MAPK RNAi oocytes, both CDK1 and Cyclin B protein levels were identical to controls (Fig. 5E). However, by Stage 14, Cyclin B levels increased dramatically, and CDK1 levels increased modestly (Fig. S8D). We analyzed the ratio of Cyclin B to CDK1 as a proxy for CDK1’s activity and assessed its change during OM. In control oocytes, CDK1 activity levels undergo minimal changes, whereas two independent MAPK RNAi lines exhibited striking increases in CDK1 activity during OM (Fig. 5F). We emphasize that increased CDK1/Cyclin B was never observed at the onset of OM, and only elevated in the Stage 14 oocyte. These findings explain the mild morphological phenotype and the normal phospho-profiles of Stage 14 oocytes depleted for MAPK. Further, the apparent increase of phosphorylated TP sites in MAPK RNAi activated eggs echoes the elevated CDK1/Cyclin B in the oocyte: by 30 minutes post-activation, CDK1 activity increases to support mitotic cycling^[5, 52]^; MAPK-depleted conditions, which contain greater starting amounts of CDK1-Cyclin B, can elicit a stronger CDK1 response in the early embryo. We note that overall protein levels appear unaffected in MAPK RNAi conditions, suggesting CDK1 and Cyclin B protein levels do not rise simply due to a general translation increase (Fig. 2E). Cyclin B translation has also been linked to PNG^[68]^. However, we did not find any premature activation of PNG in MAPK RNAi conditions, as readout by both Gnu phosphorylation and Smaug translation (Fig. S8H). We therefore conclude that CDK1 can raise its activity to compensate for reduced MAPK signaling.

CDK1 is critical for remodeling the phosphoproteome during OM and EA (Fig. 2E), but has a surprisingly mild effect on gross morphogenesis (Fig. 5C-D). This may suggest that MAPK can permit oocyte development in lieu of CDK1, and most phospho-changes driven by CDK1 are critical for releasing Prophase I arrest and/or establishing Metaphase I arrest^[31]^. In *Xenopus* and mouse, Mos signaling acts upstream of MAPK^[69, 70]^. Interestingly, in *Drosophila,* Mos is not needed for meiotic progression, but appears to act upstream of MAPK as in other systems*^[71]^*. Our results revealed that MPF levels rise in MAPK depleted conditions, possibly obscuring the normal requirements for Mos/MAPK. It will be interesting to determine whether *mos* mutant oocytes, like those lacking MAPK, similarly employ excess MPF to achieve faithful OM.

In vertebrates, CDK1 and MAPK are the prominent regulatory kinases that drive OM^[72, 73]^. In mouse oocytes, both kinases have distinct roles in promoting the degradation of securin, a crucial step following APC/C activation^[74]^. In *Xenopus*, MAPK activity stimulates MPF^[75, 76]^. Our results show that loss of MAPK, but not CDK1, hampers *Drosophila* oocyte morphogenesis; however, unlike vertebrates, loss of *Drosophila* MAPK results in increased MPF. Deactivation of MPF by overexpressing Wee1 in MAPK RNAi conditions dramatically reduced OM efficiency, suggesting that CDK1 and MAPK normally serve distinct roles, but can overlap in function to support the same processes.

Cyclin B and CDK1 levels are normal in MAPK RNAi Stage 12 oocytes and only rise in the mature Stage 14 oocytes (Fig. 5E), suggesting a temporally controlled hyperactivation of CDK1 when MAPK is depleted. These results also suggest that MAPK does not simply negatively regulate CDK1 activity, but that signaling during a critical window of oocyte development ensures adequate phosphorylation of phospho-S/T-P targets. The Stage 14-specific response to perturbed MAPK/CDK1 signaling is not limited to elevated MPF: in CDK1 RNAi or CA-Calcineurin conditions, Smaug translation is never observed during Stage 13, but occurs during Stage 14 (Fig. 3E). We posit that the Stage 14 oocyte possesses an inherent, cryptic mechanism to sense that maternal macromolecules, especially phospho-protein load, are compatible with oocyte arrest prior to EA. In some conditions (e.g., raised CDK1 activity in MAPK depleted conditions), the Stage 14 oocyte can boost phosphorylation signaling to maintain arrest, and in others (e.g., CDK1 depletion), this sensing mechanism fails and premature embryonic events leach into the oocyte.

## Conclusion and broader relevance

EA is a potential biomedical target for the advancement of assisted reproductive technologies, including improvement to pro-fertility treatments like intracytoplasmic sperm injection, or methods to limit pregnancy, like non-hormonal contraceptives. Our study first defines the requirements for a major axis of phospho-regulators leading up to and during EA (Fig. 6). MAPK and CDK1 share control of OM, and, of the two, CDK1 is especially important for restricting the onset of embryonic translation. Upon calcium rise at EA, Calcineurin dephosphorylates CDK1-MAPK targets and promotes translation. Additionally, and of great importance for targeted intervention, it is critical to understand when and how regulatory gene products behave. Through the integrated analysis of OM and EA, we delineated genes whose products are phospho-regulated during one or both transitions (Fig. 2B, 5B, SFile1). Of the 203 phospho-proteins regulated during *Drosophila* OM, 93 (45.8%) are also phospho-regulated during mouse OM (Table S3) and 177 of 203 (87.2%) have human orthologs (Table S3). Phospho-protein changes have yet to be comprehensively analyzed during mammalian EA, precluding our ability to identify orthologous phospho-changing proteins. It is noteworthy that, of the 163 phospho-regulated proteins, 141 (86.5%) have human orthologs (Table S3). Our analysis thus provides conserved, phospho-regulated candidates present for oogenic and early embryogenic development but are regulated at specific processes. Such resolution lays the groundwork for understanding the potential and efficacy of target genes for advancement of reproductive therapies. Extrapolation and comparison to vertebrate models will be an important next direction.

**Figure 6:**
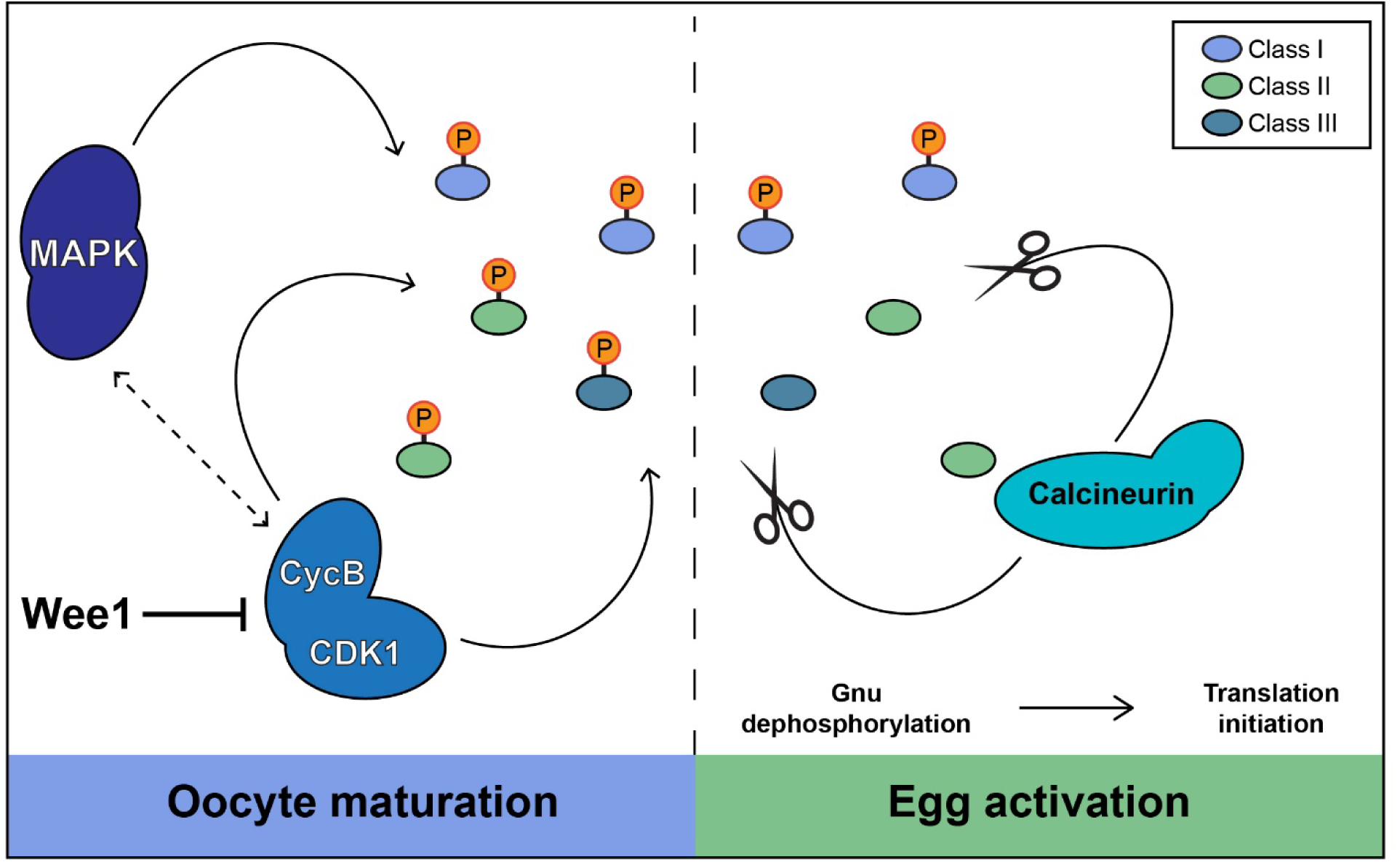
Maternal phospho-proteins primed by CDK1 and MAPK undergo Calcineurin-dependent dephosphorylation. During oocyte maturation, Class I and III maternal proteins acquire phospho-signatures via MAPK and CDK1, which promote oocyte growth and arrest before egg activation. Wee1 negatively regulates CDK1 during this time. Upon egg activation, Calcineurin dephosphorylates Class II and III substrates to facilitate the initiation of embryogenesis. Gnu is a relevant CDK1 substrate that is targeted by Calcineurin; when dephosphorylated, it complexes with other PNG members to promote the translation of Smaug.

## DATA AVAILABILITY

Mass spectrometry datasets have been deposited at the ProteomeXchange with the dataset identifier PXD087676. The code used for differential analysis and phosphorylation motif search is available on GitHub as a repository (https://github.com/WolfnerLab/JT-phosphoproteomics.git).

## ACKNOWLEDGEMENTS

We thank the Cornell Proteomics Core and BRC Imaging Facility for support in proteomics and imaging experiments. We thank Bernhard Hampoelz for the Nup358-GFP strain. We thank Masatoshi Hara and Terry Orr-Weaver for anti-Gnu antibody and Howard Lipschitz for anti-Smaug antibody. We are grateful to members of the Wolfner Lab, Anyerys Diaz, Prajal Patel, and Emily Rivard, for providing feedback on the paper. Thanks also to Arunika Das, Michael Goldberg, Marcus Kilwein, and John Schimenti for valuable feedback on the paper. We acknowledge support to WJC from grant R35GM141159 to Dr. Marcus Smolka, and grant R01HD114777 to MFW and support from a Mann award to JMT.

## AUTHOR CONTRIBUTIONS

JMT – conceptualization, data curation, formal analysis, investigation, methodology, validation, visualization, writing – original draft, writing – review & editing. WJC, methodology, formal analysis. SD, methodology, writing – review & editing. MFW, conceptualization, resources, supervision, funding acquisition, writing – original draft, writing – review & editing.

## Abbreviations

EA: egg activation
OM: oocyte maturation;
CDK1: Cyclin dependent kinase 1;
MPF: maturation promoting factor;
MAPK: mitogen activated protein kinase;
PNG: Pan Gu Kinase

---

**Figure S1:**
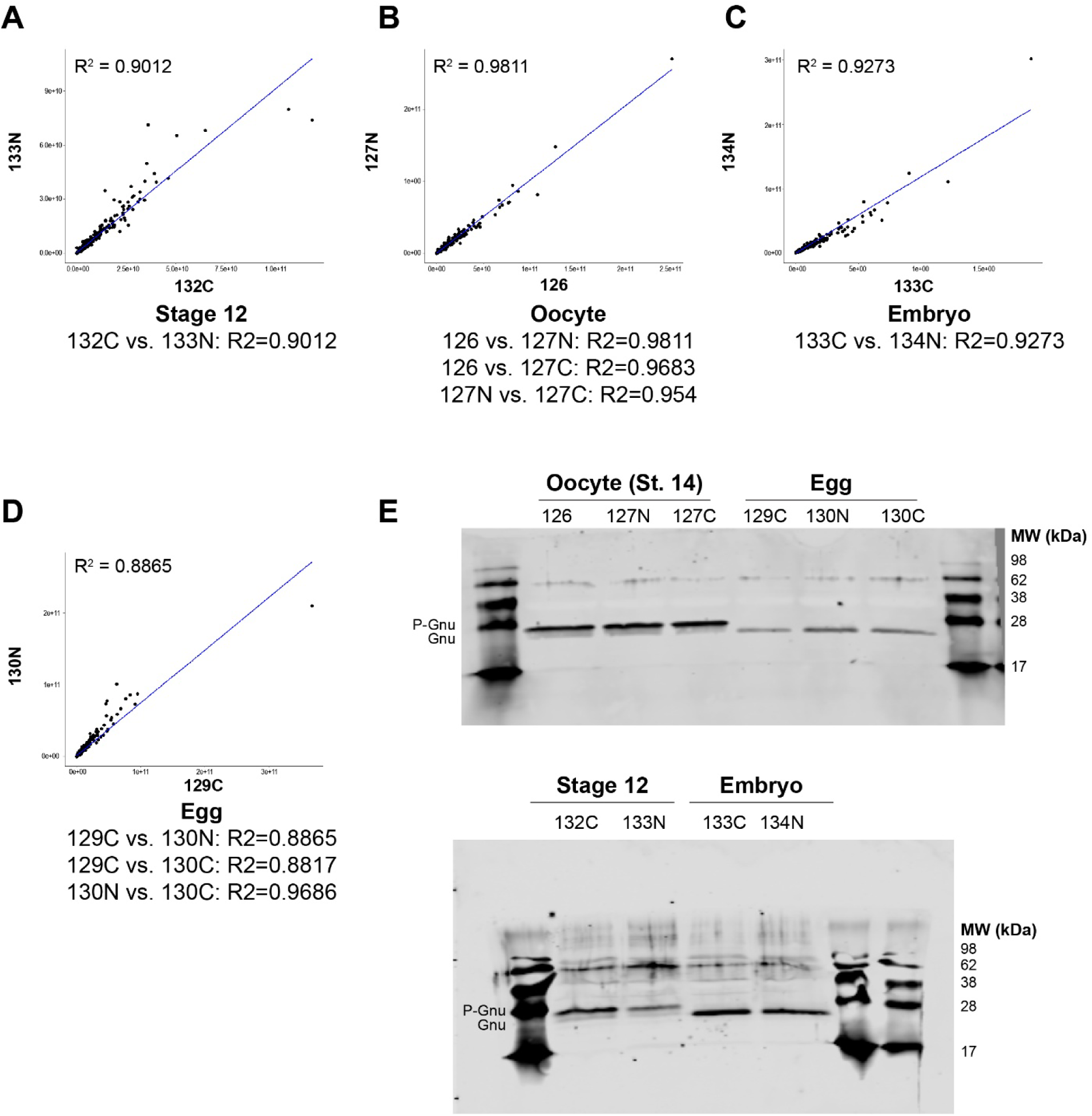
Rigor of phospho-LC-MS/MS sample preparation. A-D) For each sample type included in the TMT-multiplex, we performed simple linear regression between each sample to determine how comparable replicates were. Each comparison yielded an R^2^ value greater than 0.88, indicating high correlation between independently prepared replicates. E) Protein lysates used for phospho-LC-MS/MS were analyzed for Gnu phospho-state, which is mostly phosphorylated in Stage 12 and 14 oocytes, but dephosphorylated in Eggs and Embryos.

**Figure S2:**
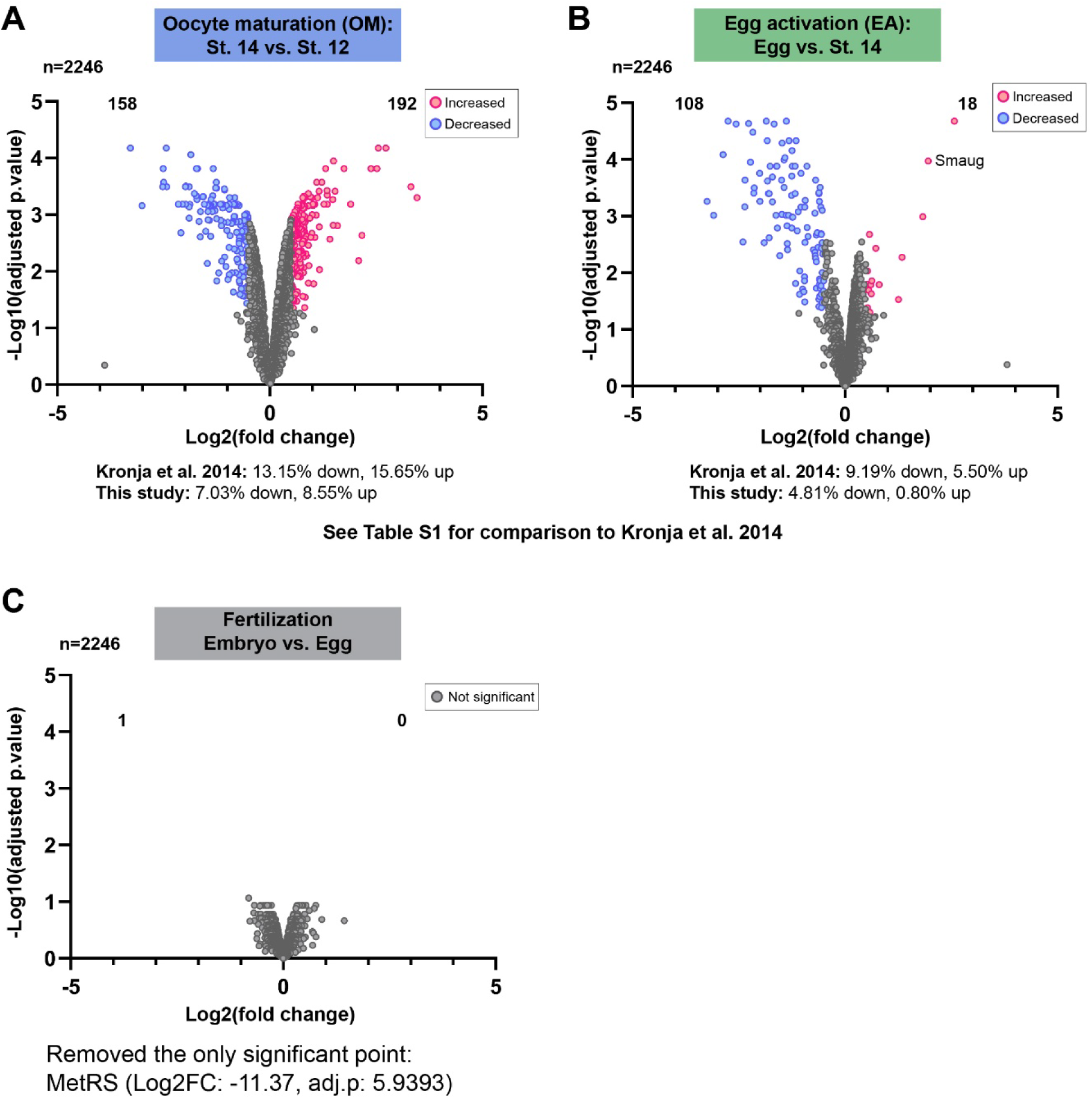
Global proteome analysis of oocyte maturation, egg activation, and fertilization. A) During OM, 350 of 2246 (15.58%) proteins undergo abundance changes. B) During EA, 126 of 2246 (5.61%) proteins change significantly. We compared these trends to a former proteome analysis of both transitions (Kronja et al. 2014; Table S1) and found similar trends, whereby there are stronger protein increases during OM and stronger decreases during EA. C) The proteomes of activated fertilized and unfertilized eggs were nearly identical. A single change to MetRS was observed. For all analyses shown, pairwise comparisons were performed using Bayes moderated t-tests via LIMMA. Significant changes were those with a Log2 fold change >+/-0.5 and adjusted p value <0.05.

**Figure S3:**
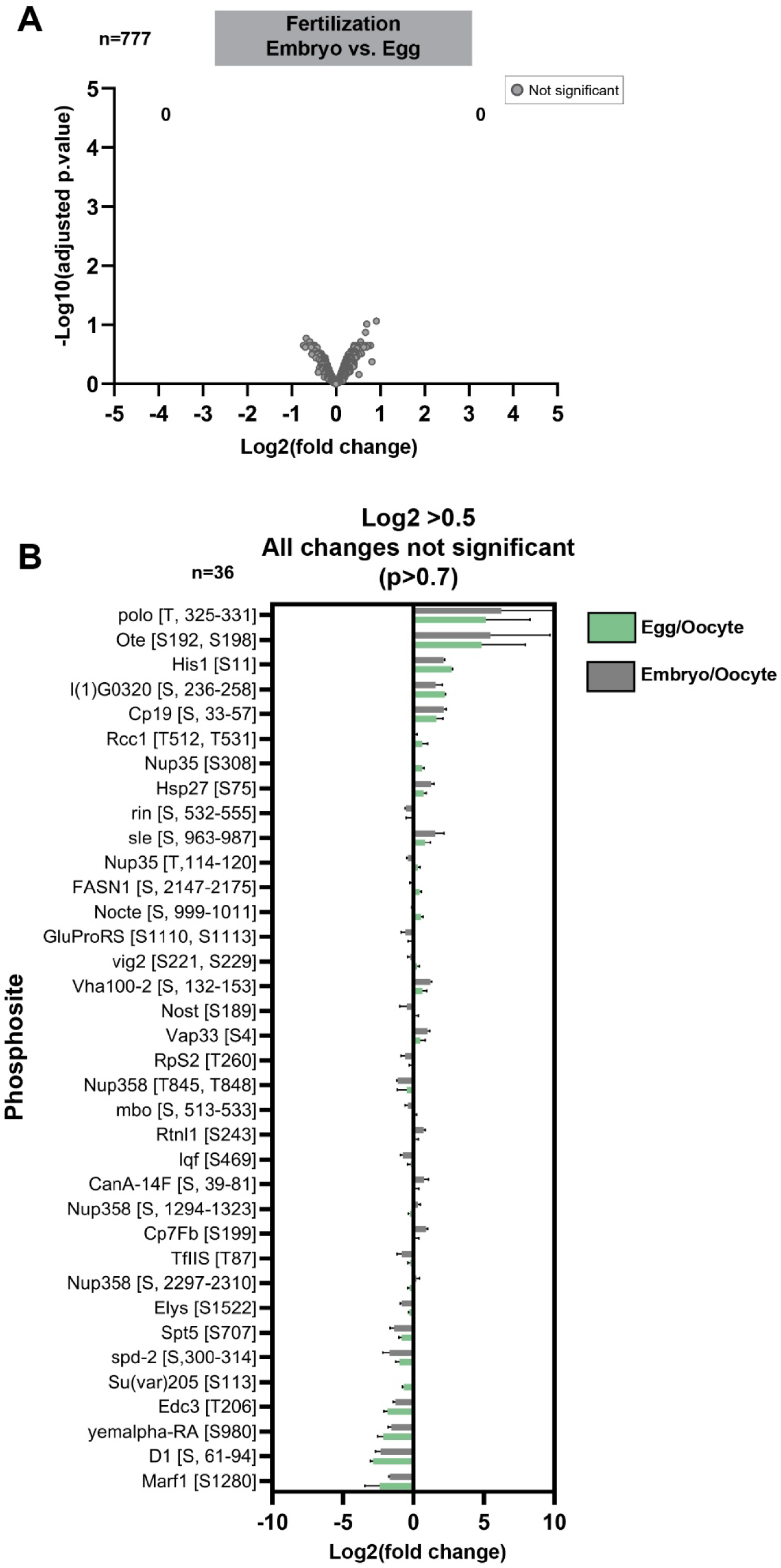
Phosphorylation analysis of fertilization compared to egg activation. A) Phospho-protein abundance was compared between activated fertilized embryos and unfertilized eggs of the equivalent timepoint post-egg laying (<30 minutes). B) Phospho-protein Log2 fold changes between Egg/Oocyte and Embryo/Oocyte, whereby differential analysis between them showed no significant differences. Phosphopeptides with a difference of >+/-0.5 between Egg/Oocyte and Embryo/Oocyte comparisons are displayed; however, we note that all p-values are greater than 0.7 and these changes are not significant.

**Figure S4:**
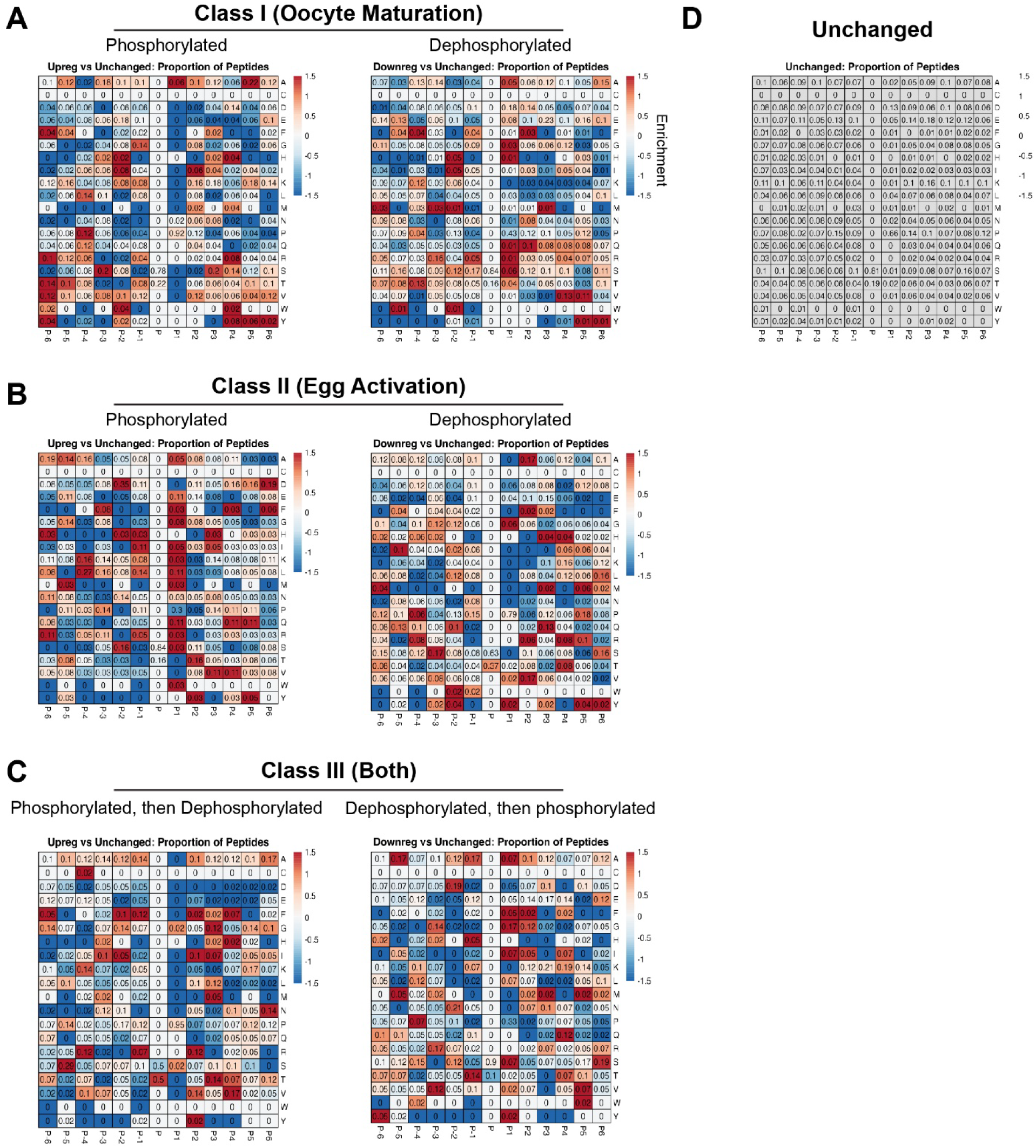
Unbiased phospho-motif enrichment analysis. A-C) Heatmaps generated by phospho-motif enrichment analysis for each class of phospho-sites. Along the x axis, position ‘P’ represents the site of phosphorylation and P-6 through P6 represent the positions of flanking amino acids. On the y axis, each amino acid is displayed. Numbers represent proportion of each amino acid’s occurrence in positions P-6 through P6. Inputs for each comparison were phosphopeptides identified as significantly changing by our differential analysis for the conditions shown. To determine enrichment, each class of phospho-sites was compared to the statistically unchanged (background) pool of phospho-sites from our analysis (D). Red indicates overrepresentation and blue indicates underrepresentation.

**Figure S5:**
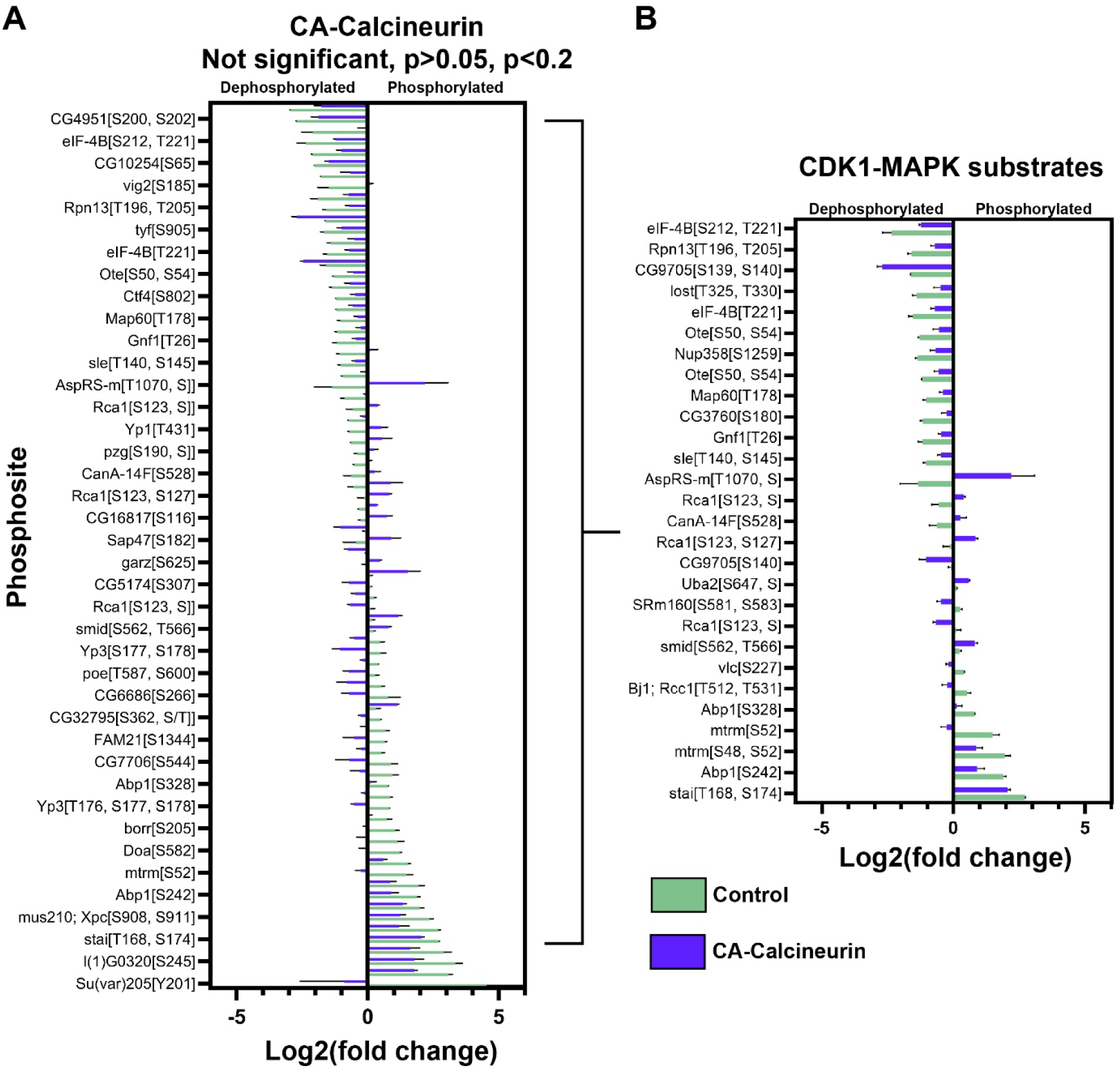
Reanalysis of CA-Calcineurin phospho-sites from Zhang et al. 2019. By reanalyzing Zhang et al. 2019, we found no significant changes to phospho-proteins when CA-Calcineurin is expressed compared to controls. However, we found 80 phospho-sites that trended towards significance at p<0.2 (A). Among these were 28 CDK1-substrates (B). Log2 fold changes of Egg/Oocyte between control and CA-Calcineurin were compared via Bayes moderated t-tests.

**Figure S6:**
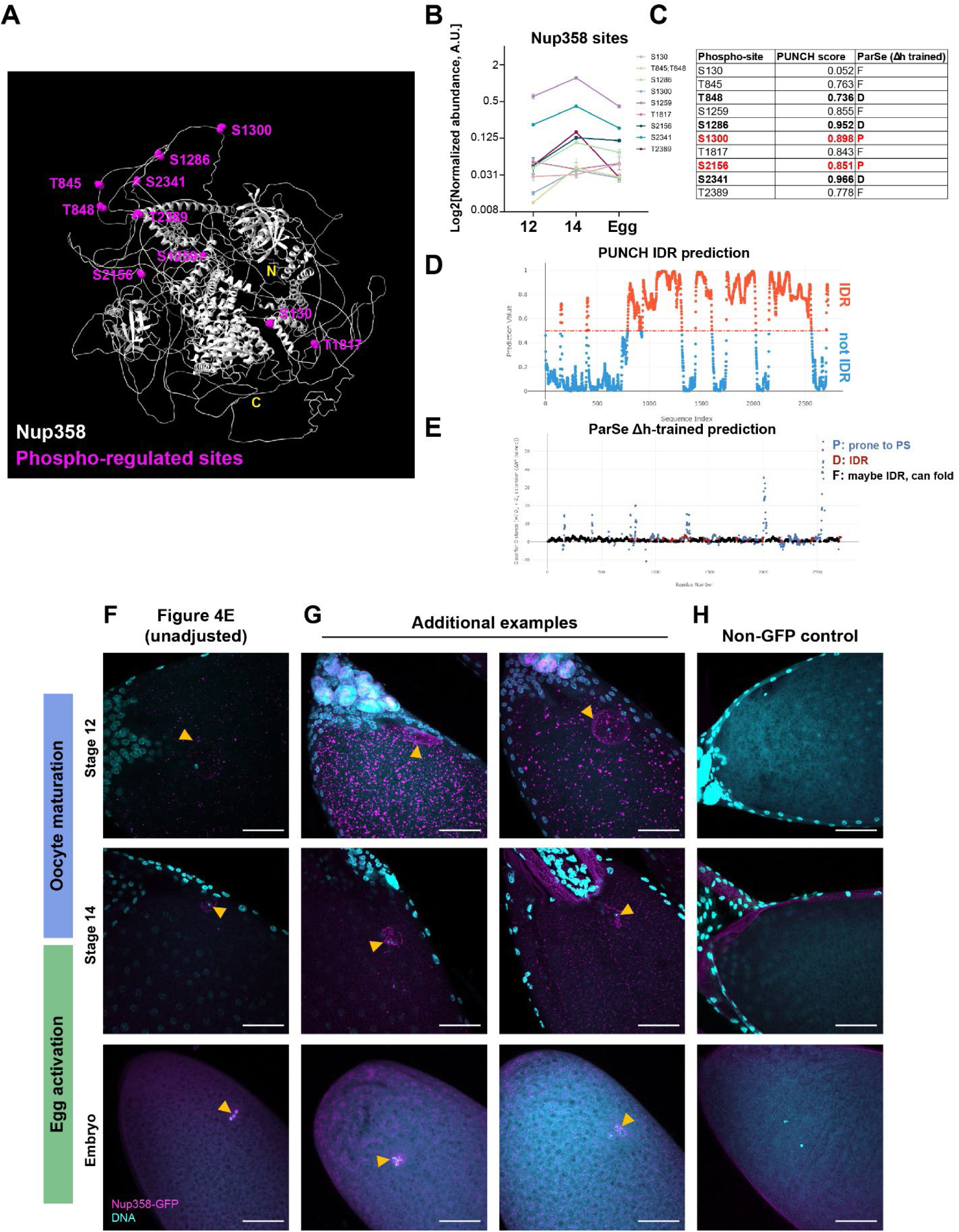
Analysis of Nup358. A) Alphafold2 predicted structure of Nup358 with all phospho-regulated sites mapped (magenta). B) Abundance traces for all phospho-regulated Nup358 sites. C-E) Outputs from PUNCH and ParSe analyses. PUNCH scores greater than 0.5 are predicted to be intrinsically disordered regions (IDRs). Bolded sites in C) are predicted as IDRs by both algorithms. Red sites are predicted to be prone to phase separation (PS) by ParSe analysis. P: sites prone to phase separation. D: IDR. F: Sites that may be IDRs, but can likely fold. F) Unedited micrographs from Fig. 3E. G) Additional examples of Nup358’s localization to the GV (Stage 12), apposition to the metaphase I spindle (Stage 14), and presence on polar bodies or polar body aggregates. Relevant structures are marked with yellow arrowheads. H) Wild type, non-GFP expressing controls for each stage. Scale bars = 50μm.

**Figure S7:**
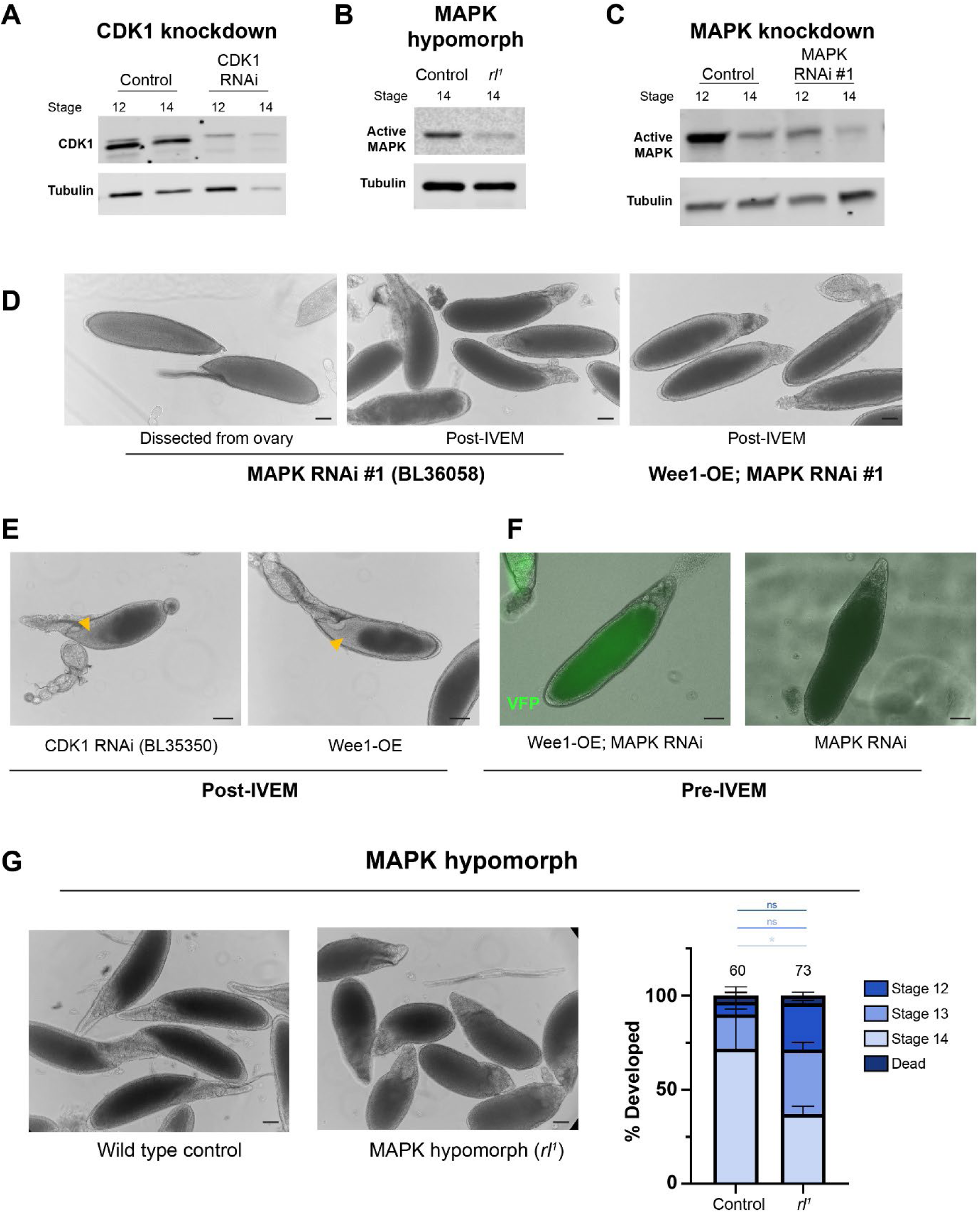
Supporting information for IVEM assay. A-C) Western blot validation of reduced CDK1 levels (A) or MAPK activity (phospho-MAPK, B-C) in indicated knockdown or hypomorph conditions. CDK1/MAPK reduction was confirmed at least three times per condition. D) Examples of MAPK RNAi oocytes dissected from ovaries or matured in culture. In both conditions, dorsal appendage formation is stunted. Additional examples of Wee1-OE;MAPK RNAi oocytes matured in culture are shown. E) Examples of CDK1 RNAi and Wee1-OE oocytes after IVEM which display a clear region of ooplasm (orange arrowhead). F) Validation of Wee1-OE (UASp-Wee1-VFP crossed to Maternal alpha-Tubulin Gal4) by fluorescence imaging of VFP signal. VFP was present in all Wee1-OE samples (shown: Wee1-OE;MAPK RNAi) but absent from non-VFP expressing samples. G) IVEM of MAPK hypomorph oocytes (*rl^1^*). Compared to controls, *rl^1^* oocytes showed a significant reduction in their capacity to reach Stage 14 in culture. All scale bars = 50μm.

**Figure S8:**
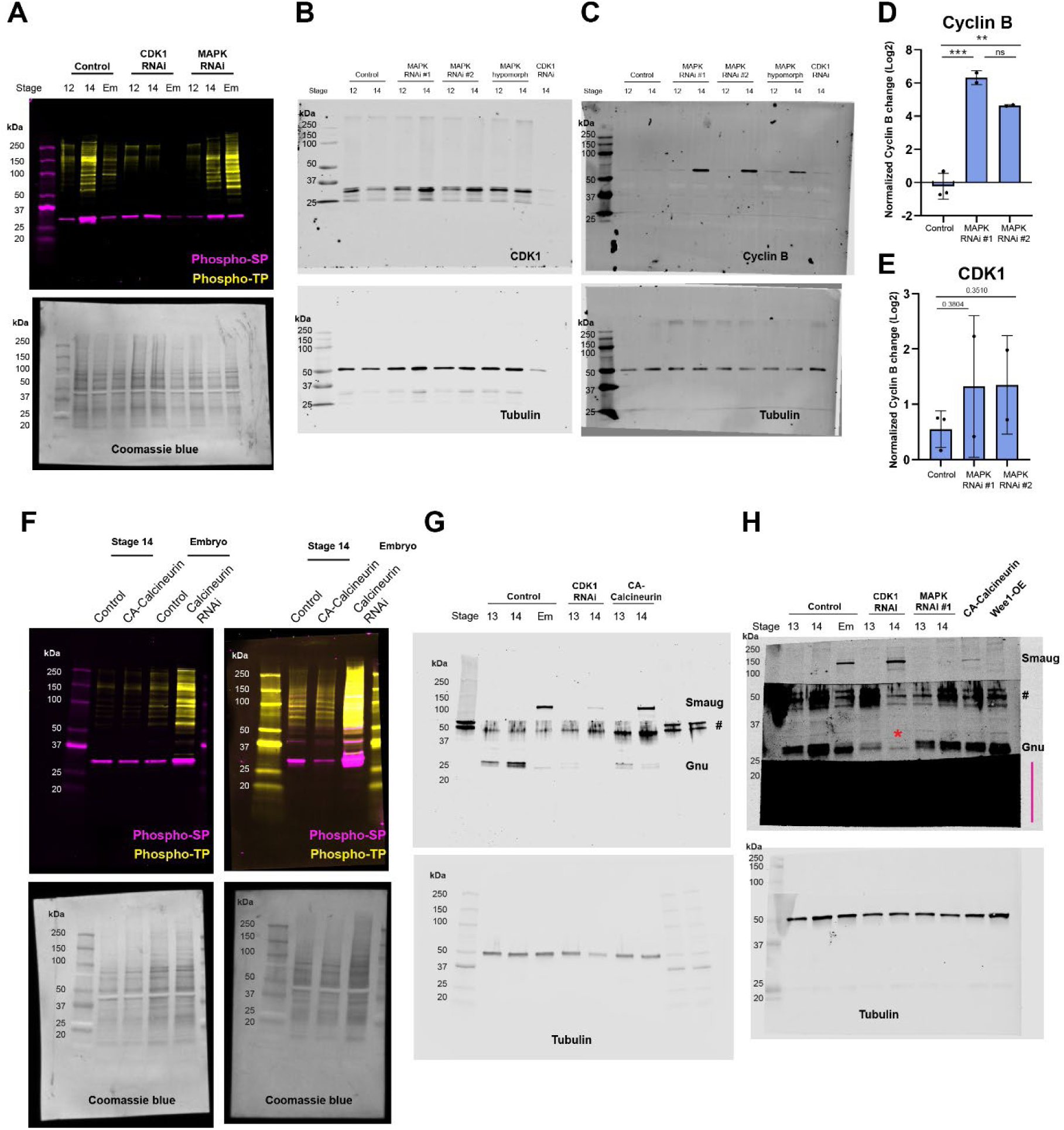
Full western blots. Full scans for all western blots. A) Full blot and Coomassie stain for Fig. 2E. B-E) Full blots of anti-CDK1, anti-Cyclin B, and anti-Tubulin for Fig. 5E. Quantification of Cyclin B and CDK1 levels are shown individually; ratios are analyzed in Fig. 5F. F) Full blots and Coomassie stains for Fig. 3A. G) Full blots for Fig. 3E. H) Additional example of data shown in Fig. 5E-H, whereby the blot is exposed to show the presence of dephosphorylated Gnu in CDK1 RNAi oocytes (red asterisk). The region indicated by a magenta line is probed for an antibody not included in this study. # indicates nonspecific bands from anti-Gnu antibody.

## Notes

### Competing Interest Statement

The authors have declared no competing interest.

